# Pseudouridine selects RNAs for extracellular transport

**DOI:** 10.1101/2025.10.28.685156

**Authors:** Alessandro Scacchetti, Tuan D. Tran, Emily J. Shields, Lauren N. Reich, John F. Doherty, Julia A. Tasca, Grace E. Lee, Alejandro A. Vilcaes, Richard Lauman, Natali L. Chanaday, Benjamin A. Garcia, Colin C. Conine, Roberto Bonasio

## Abstract

RNAs move through the extracellular space to transmit information between cells, including mammalian neurons, yet how specific RNAs are channeled into these extracellular routes is unknown. Using genome-wide CRISPR screening, proteomics, and high-sensitivity transcriptomics in a neuronal model system, we identify domesticated retroviral proteins and RNA-modifying enzymes that regulate RNA loading into and transportation via extracellular vesicles. We show that the pseudouridine synthase PUS1 is a key determinant of RNA trafficking, and that its catalytic product in RNA, pseudouridine, is enriched in extracellular RNAs from transformed and primary neurons. Furthermore, the presence of pseudouridine on select RNAs is both necessary and sufficient for their extracellular export. Finally, we show that myosin light chain 6 (MYL6) is a pseudouridine-binding protein required for secretion of synthetic and endogenous RNAs. These findings reveal a biochemical code linking chemical RNA modification to extracellular transport, and establish a framework to study the function of extracellular RNAs in the nervous system and beyond.

## INTRODUCTION

Cells exchange information to coordinate their actions. In addition to hormones, neurotransmitters, and other small molecules, RNAs also cross the plasma membrane and traverse the extracellular space to exert regulatory functions across cellular, tissue, and sometimes organismal boundaries^1–6^. In mammals, such intercellular transfer of RNA has been associated with diverse physiological and pathological processes, including brain function and disease^7–10^, cancer^11^ and intergenerational epigenetic inheritance^12^, but its mechanistic bases remain obscure. Here, we investigate how RNAs are selected for extracellular export, a necessary step toward understanding what type(s) of information they communicate between cells, and determining whether and how this process affect phenotypic, transcriptional, and epigenetic states.

Extracellular RNAs (exRNAs) are often found within extracellular vesicles (EVs), membrane-based particles of heterogenous size, origin, and composition^13,14^, produced by most cell types^15^, and capable of transferring their cargo between cells^16^. Other structures, including membrane-free ribonucleoprotein complexes^17,18^ and virus-like particles^19–23^, can also transport RNA through the extracellular environment, although these processes are even less well understood than those mediated by canonical EVs.

The exRNAs produced by a given cell type do not recapitulate its intracellular RNA content^24–28^, suggesting the existence of molecular pathways that facilitate (select) or prevent (filter) the release of specific RNAs in the extracellular environment. Mechanisms for exRNA selection have been proposed, including sequence-specific sorting mediated by RNA-binding proteins^27,29–33^ and selective processing of exRNAs after their release^34–36^. To date, mechanistic studies have focused on extracellular transport mechanisms for microRNAs (miRNAs), largely in non-neuronal cells. However, miRNAs are relatively scarce in EVs^37^, and it is becoming increasingly clear that neurons might have specialized strategies to select and release exRNAs^19,20,23,38^. Hence, the full spectrum of mechanisms that participate in exRNAs selection and transport, especially in neurons, remain unknown.

Here, we report a novel genetic screen that uncovered new regulators of exRNA export in a neuronal cell line. These include domesticated retroviral proteins and RNA-modifying enzymes. Among the latter, we show that the pseudouridine synthetase 1 (PUS1) and its catalytic product, pseudouridine (Ψ), are necessary and sufficient for the selection and extracellular release of certain exRNAs, including full-length transfer RNAs (tRNAs) and small nucleolar RNAs (snoRNAs), through a process that also requires the non-canonical pseudouridine binding protein myosin light chain 6 (MYL6).

### Neuronal CADs produce RNA-containing EVs

To investigate the genetic and biochemical determinants of neuronal exRNA biogenesis in a tractable model, we selected Cath.a-differentiated cells (CADs), an immortalized cell line that resembles catecholaminergic neurons and is capable of forming neurite-like structures under serum-free conditions^39–41^. We isolated EVs from conditioned, serum-free medium from CAD cultures using an established serial ultracentrifugation method (**Fig. 1A**)^42^. This relatively crude EV fraction consisted primarily of particles ranging from 90 to 150 nm in diameter (median 123 nm, **Fig. 1B**), consistent with the expected size range of small EVs^13^. This fraction was enriched for known EV markers (SDCBP, PDCD6IP/ALIX, TSG101, CD9), and depleted for conventional control mitochondrial and ER proteins (COXB5 and CANX) (**Fig. 1C**)^43^. Overall, 81% of the proteins enriched in our EV isolate also appeared in a curated database of EV components (Vesiclepedia, **Fig. S1A**)^44^, and gene ontology (GO) annotations for the proteins we identified in CAD EVs were enriched for relevant terms, including “*intracellular vesicle*” and “*endosome*” (**Fig. S1B**). Consistent with the neuronal identity of CADs, GO terms associated with neurons (“*synaptic membrane*”, “*axon*”) were also enriched, as well as specific neuronal EV markers L1CAM and ATP1A3^45^ (**Fig. 1C**, **Table S1**).

**Figure 1.**
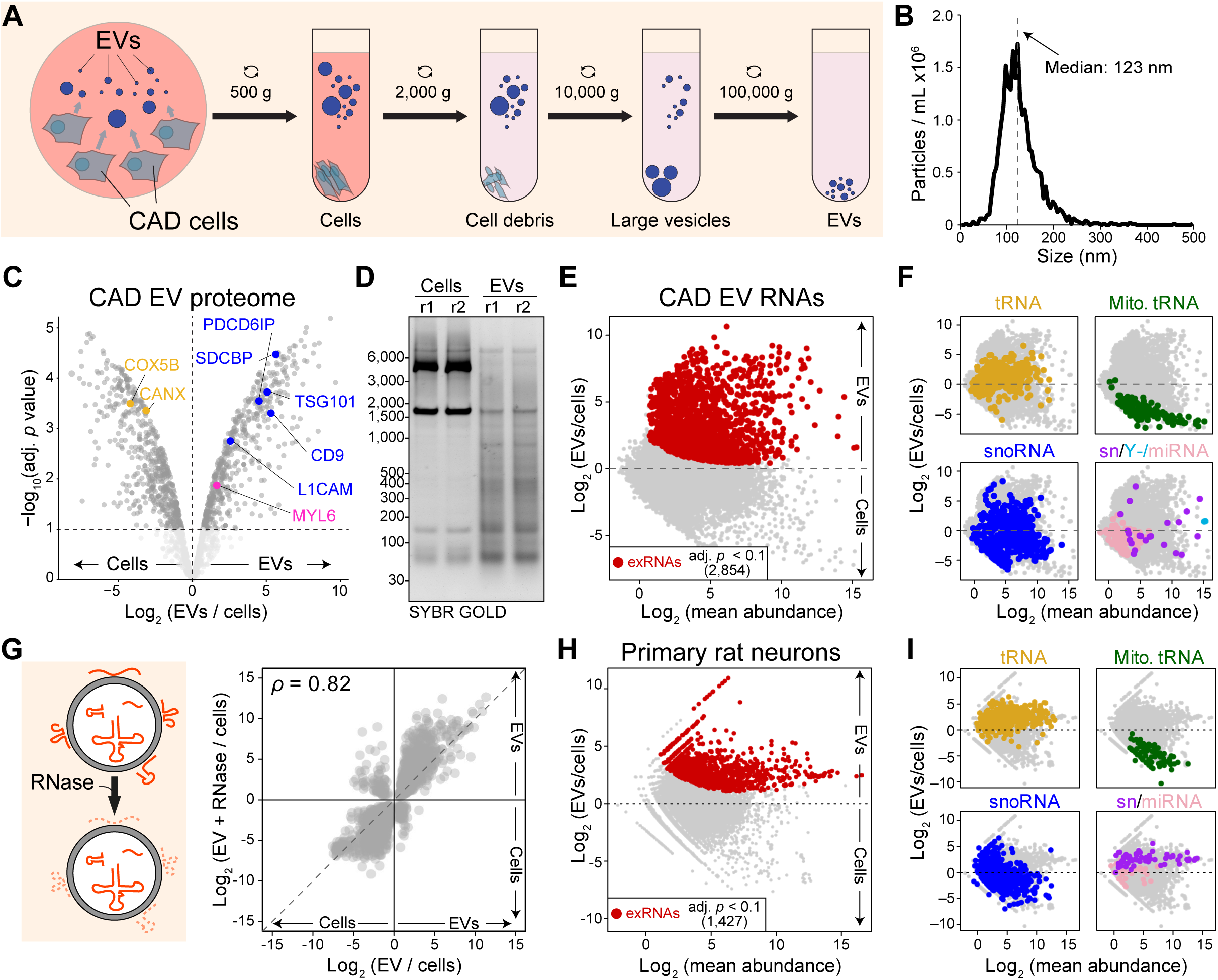
CADs and primary neurons produce EVs enriched in tRNAs and snoRNAs. (A) Scheme of EV purification protocol. Conditioned medium was collected and subjected to several round of centrifugation at increasing speeds to remove cells, cell debris, and large vesicles. In the last step EVs were pelleted via ultracentrifugation at 100,000 x *g*. (B) Nanoparticle tracking analysis of EVs from CADs. The plot shows the average (*n* = 2) number of particles per mL on the *y* axis and their size on the *x* axis. The vertical dashed line indicates the median size. (C) Comparison of cellular and EV proteomes by mass spectrometry. Gray, significantly different proteins (adjusted *p* < 0.1; *n* = 2); blue, EV markers; yellow, cellular markers. (D) Gel electrophoresis (2% agarose-formaldehyde) comparing cellular *vs.* EV RNA. Individual replicates were loaded in separate lanes. (E) RNA content of CADs EVs *vs.* cells, measured by deep sequencing (LIDAR). Red, transcripts significantly enriched in EVs [exRNAs; adjusted *p* < 0.1 and log_2_(EVs/cells) > 0; *n* ≥ 3]. (F) Same as (E) with different classes of RNA highlighted by colors, as indicated. (G) Left: EVs were incubated with RNase A/T1 to digest all RNAs not protected by a lipid bilayer. Right: enrichment of exRNAs (EVs vs. cells) in EVs from CADs before (x axis) and after (y axis) RNase treatment. Only exRNAs significantly different (adjusted *p* < 0.1) in either treatment are shown. *ρ*, Spearman’s correlation value. (H–I) Same as (E–F) but for EVs purified by size-exclusion chromatography from cultured rat primary neurons (*n* = 2).

In contrast to the intracellular RNA composition of CADs, their EVs contained predominantly small RNA species (< 500 nts; **Fig. 1D**), as previously observed in other neuronal EVs^46^. Because conventional, ligation-based protocols to sequence small RNAs were largely developed to detect miRNAs^47,48^, we used LIDAR sequencing^49^, which captures a broader range of RNAs regardless of their size and chemical modifications, to obtain a comprehensive profile of the exRNAs produced by CADs. LIDAR revealed 2,854 RNAs that were significantly (adjusted *p* < 0.1) enriched in EVs compared to whole-cell extract (**Fig. 1E**). Compared to the intracellular transcriptome, CAD exRNAs were enriched for nuclear tRNAs, Y-RNAs, and some small nuclear RNAs (snRNAs), and depleted for mitochondrial tRNAs and miRNAs (**Fig. 1F; S1C**). Some classes of snoRNAs such as the H/ACA box family were enriched in CAD EVs, whereas others, such as C/D box snoRNAs were more abundant inside the cells (**Fig. S1D**). Further inspection of the reads mapping to tRNA genes revealed the presence in EVs of a mixture of full-length tRNAs and various types of tRNA-derived fragments (tDRs) (**Fig. S1E–F**).

To confirm that the exRNAs produced by CAD cells resided in the lumen of EVs, we exposed the crude EV isolate to a mixture of RNase A and T1. A substantial amount of exRNAs resisted RNase treatment in absence but not in presence of a detergent (Triton X-100), indicating that these exRNAs resided in the lumen of membrane-delimited EVs (**Fig. S1G**). LIDAR sequencing after RNase treatment of the EV isolate confirmed that most of the exRNAs we detected were not simply associated with EVs, but were loaded into EVs as cargo and thus protected from degradation by the EV membrane (**Fig. 1G**).

To extend our observation to *bona fide* neurons, we isolated EVs from cultured rat primary cortical neurons, using a size-exclusion chromatography protocol^50^, which yielded particles of size comparable to those obtained from CADs (**Fig. S1H**). LIDAR sequencing of the RNA content of these EVs revealed an enrichment for specific classes of RNAs, remarkably similar to those found in CAD EVs (**Fig. 1H–I**), suggesting the existence of conserved mechanisms of neuronal exRNA selection.

Thus, neuron-like CAD cells produce EVs that contain known EV-associated and neuronal proteins, and are enriched in specific classes of RNA (including tRNAs and their fragments), similar to EVs produced by primary neurons.

### Single-guide RNAs are loaded into EVs

The fact that CAD EVs are enriched for specific RNAs when compared to their intracellular content suggests the presence of dedicated pathways ensuring that only certain transcripts are selected to become exRNAs. Potential mechanisms for both active and passive regulation of RNA secretion have been proposed^51^, but very few examples, seemingly dedicated to the export of specific miRNAs, have been characterized^27,30,31,52^. It is unclear whether these bespoke pathways also affect other RNA classes, which constitute the majority of exRNAs (**Fig. S1C**).

In search for general regulators of exRNA biogenesis, we performed a genome-wide knockout screen. In a typical CRISPR/Cas9 screen, a library of single-guide RNAs (sgRNAs) targeting genes of interest (or all genes in the case of genome-wide screens) is transduced into cells that are then selected for a phenotype, for example drug resistance. Enriched or depleted sgRNAs provide information on genes that regulate that phenotype and are identified by sequencing the encoding transgenes integrated into the genome of the selected cells, thus linking genotypes to phenotypes. This approach cannot be used for phenotypes not physically connected to the cell of origin, such as EV abundance and composition.

We made the serendipitous observation that sgRNAs expressed from a lentiviral vector could be detected in the crude EV fraction from CADs (**Fig. S2A**), therefore coupling the sgRNA genotype of the originating cell to phenotypes associated with the sgRNA-containing EVs (**Fig. 2A, S2B**). We reasoned that knockout of genes required for EV production or for loading RNAs into EVs would result in the disappearance of the corresponding sgRNAs from the EV fraction, allowing us to perform a genome-wide screen to identify genes involved in these processes.

**Figure 2.**
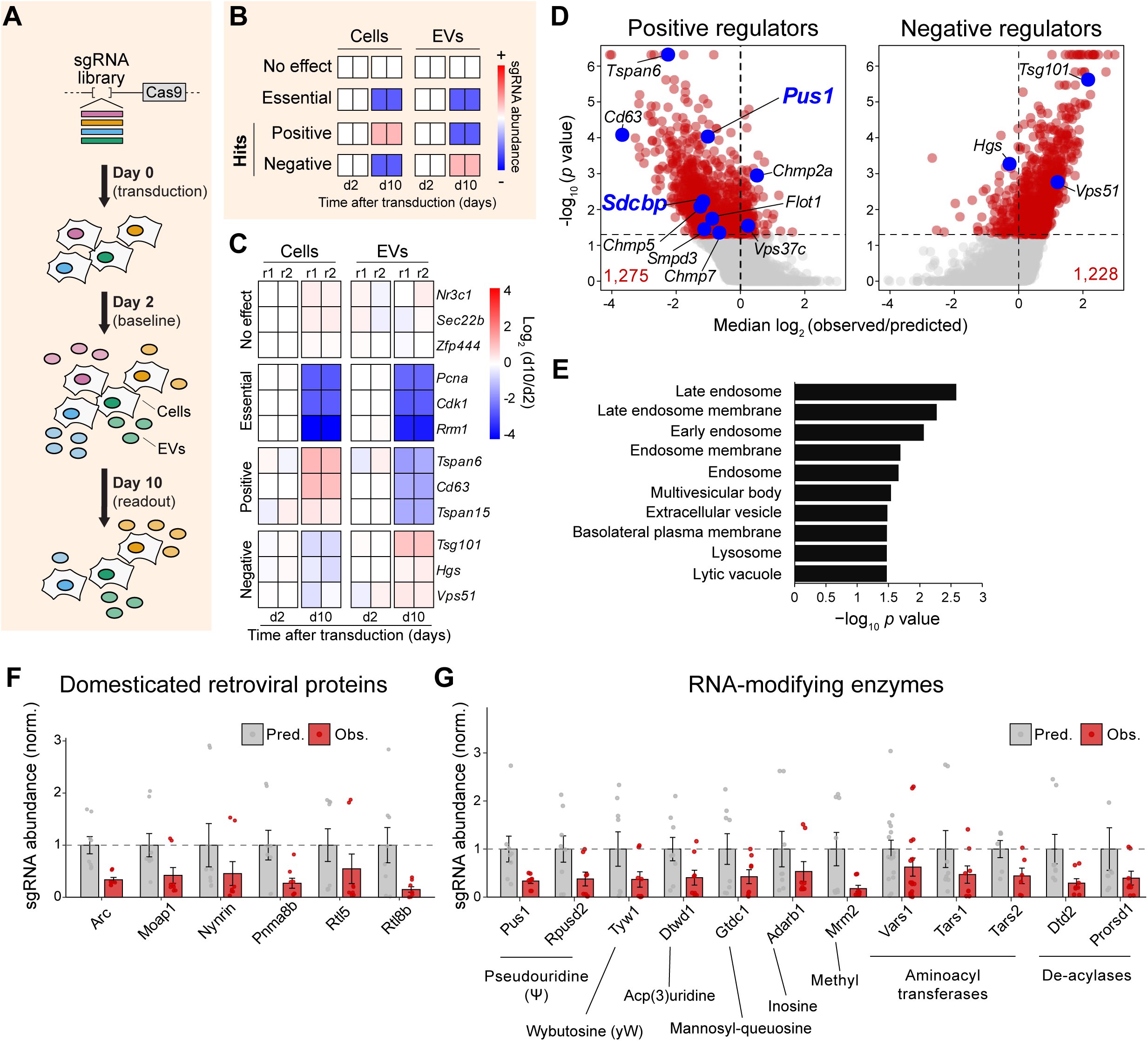
A genome-wide screen identifies regulators of RNA secretion. (A) Scheme of the genetic screen. CADs were transduced with a lentiviral library containing sgRNAs against most protein coding genes. Two (baseline) and 10 (readout) days later, cells and EVs were harvested to measure sgRNA levels. (B) Hypothetical gene categories with expected changes in sgRNA abundance over time in cells and EVs. (C) Change in sgRNA abundance (*n* = 4 sgRNAs per gene), compared to their average abundance in the corresponding sample (cells or EVs) at day 2. Individual biological replicates (r1 and r2) are shown. (D) Observed/predicted ratio of sgRNAs targeting putative positive (left) and negative (right) regulators of RNA secretion, derived from MaGECK analysis. Significant hits (*p* < 0.05) are in red (*n* = 2). Candidates mentioned in the text are highlighted in blue. (E) GO analysis (cellular component) of significant positive regulators from (D). (F) Predicted (gray) vs. observed (red) sgRNA abundances in EVs (± SEM), normalized to the average predicted abundance (see methods), for retrovirus-derived proteins identified as significant (*p <* 0.05) positive regulators from MaGECK analysis. Values for individual sgRNAs are shown. (G) Same as (F) but for RNA-modifying enzymes.

We envisioned that targeted genes and the corresponding sgRNAs would fall within 4 possible categories (**Fig. 2B**): 1) sgRNAs not affecting viability, EV production, or RNA export (“no effect”) would remain constant over time in both cells and EVs; 2) sgRNAs targeting genes promoting survival or proliferation would be depleted over time in both intracellular and extracellular pools (“essential”); 3) positive regulators of EV production or RNA export would become less abundant in the EV-associated pool over time, while possibly accumulating within cells (“positive”); and 4) *vice versa* for negative regulators (“negative”).

### A genome-wide screen for EV regulators

We transduced CADs with the genome-wide Brie library, comprising ∼80,000 sgRNAs targeting ∼20,000 protein-coding genes (**Fig. 2A**)^53^. Two days after infection, we harvested a fraction of the cells and EVs to quantify baseline sgRNA distribution. Eight days later (day 10 after transduction), we harvested cells and EVs again, and prepared libraries to sequence the sgRNAs, using a modified LIDAR protocol (**Fig. S2C**). We performed the screen twice, obtaining two independent experimental replicates.

Deep sequencing detected > 95% of sgRNAs in all samples (**Fig. S2D**). As early as day 2 after transduction, likely before most knockout effects would manifest, not all sgRNAs were equally represented in EVs compared to cells (**Fig. S2E**), indicating that some sgRNAs are more efficiently loaded into EVs than others. This suggests that these exogenous RNAs are likely subjected to the same selection processes that act on endogenous RNAs, potentially influenced by differences in sequence, structure, or both. Consistent with this, sgRNAs that were preferentially exported as exRNAs were characterized by a higher frequency of uridines at the 5’ (**Fig. S2F**), raising the possibility that these uridines, or perhaps their chemical modification, might contribute to the selection process.

We analyzed the relative changes in abundance of sgRNAs in cells and EVs over time and sorted the sgRNAs in the four categories described above (**Fig. 2C**). As expected, most sgRNAs were neither enriched nor depleted (“no effect”), while sgRNAs targeting reported essential genes, such as *Pcna*, *Cdk1*, and *Rrm1* were depleted at day 10 compared to day 2, both in cells and in EVs (“essential”). Single-guide RNAs targeting tetraspanins, *Cd63*, *Tspan6*, and *Tspan15* were depleted at day 10 in EVs, but not inside the cells, where they were either unchanged or enriched. These genes might function as positive regulators of EV biogenesis (“positive”), in line with their known role in membrane trafficking^54^. Conversely, we observed an enrichment of sgRNAs in EVs, but not in cells, (“negative”) for two subunits of ESCRT complexes, *Tsg101* and *Hgs* (*Hrs*), consistent with recent reports suggesting that they may function as negative regulators for release of some EV subtypes^55,56^. *Vps51*, encoding a subunit of the GARP complex^57^, was also classified by our screen as a negative regulator, likely due to its role in mediating retrograde trafficking from early endosomes to the Golgi apparatus^58^.

For each sgRNA, we calculated a “predicted” abundance in EVs compared to cells at day 10 based on its export efficiency measured at day 2 and then determined deviations from this predicted value in the “observed” sgRNA abundances. Overall, we identified 1,275 and 1,228 genes targeted by significantly (*p* value < 0.05) depleted or enriched sgRNAs (**Fig. 2D; Table S2**), and therefore likely to function as positive or negative regulators of exRNA selection/transport, respectively. Overall, the candidate positive regulators in our screen were enriched for GO terms related to known EV biogenesis routes, such as “*late endosome*”, “*multivesicular body*”, and “*extracellular vesicle*” (**Fig. 2E**), and comprised various genes involved in the regulation of EV production and release (*Tspan6*, *Cd63*, *Flot1*, *Smpd3*, *Vps37c*, *Chmp2a*, *Chmp5*, and *Chmp7*) (**Fig. 2D**). Among these, *Sdcbp* (**Fig. 2D; S2G**) stood out because of its demonstrated role in EV biogenesis^59^, established use as marker for these structures^43^, and abundance in CAD EVs (**Fig. 1C**). To confirm its function in neuronal EVs, we generated two independent *Sdcbp^-/-^* CAD clones (**Fig. S2H**). Loss of SDCBP caused reduced particle numbers indicative of impaired EV production, consistent with the effects observed in other cell types^59^ (**Fig. S2I–J**).

Given that our screen utilized extracellular sgRNAs directly as a readout, the identification of *Sdcbp* suggests that sgRNAs are exported, at least in part, via EVs originating from a variation of the ESCRT pathway^59^. However, several genes encoding domesticated retroviral proteins (*Arc, Moap1, Nynrin, Pnma8b, Rtl5, Rtl8b*)^60^ were also among the top candidates for positive regulators of sgRNA export (**Fig. 2F**). Encapsulation of RNAs in capsid-like structures could therefore represent an alternative route for exRNA export, potentially one reserved for neuronal cells^19,20,61–63^, given that several of these genes are preferentially expressed in the nervous system^60^. Consistent with neuronal-biased expression, all these candidates except *Moap1* (for which our RNA-seq was not sufficiently sensitive) were upregulated upon differentiation of CADs into a more mature neuronal phenotype (**Fig. S2K**)^39,40^.

Our screen also identified positive regulators with no obvious link to membrane trafficking. We reasoned that some of these factors might be involved in RNA selection or loading into the EVs and that, therefore, their depletion would cause the disappearance of the corresponding sgRNAs over time. Amongst them, we noted several enzymes responsible for installing or removing chemical modifications on RNA, including pseudouridine (*Pus1, Rpusd2*)^64^, wybutosine (*Tyw1*)^65^, acp(3)uridine (acp^3^U) (*Dtwd1*) ^66^, mannosyl-queuosine (*Gtdc1*)^67^, inosine (*Adarb1*)^68^, methyl groups (*Mrm2*) ^69^, and amino acids (*Vars1, Tars1, Tars2, Dtd2, Prorsd1*)^70–72^ (**Fig. 2G**). More broadly, we identified 224 RNA-binding proteins amongst the positive exRNA regulators via RBP2GO analysis^73^ (**Table S3**), in line with our expectations that factors involved in the selection and loading of RNAs into EVs would be revealed by our genetic screen.

In summary, we devised a pooled genome-wide strategy to link genetic perturbation to regulation of exRNA export. We identified several factors that could be promoting or restricting exRNA selection and transport, including several domesticated retroviral genes, suggesting an alternative, potentially neuron-specific mode of RNA secretion. Our screen also uncovered a variety of RNA modifiers and RNA-binding proteins that could be involved in marking and selecting specific RNAs for extracellular trafficking, as demonstrated below for PUS1.

### PUS1 shapes the exRNA population

Among the RNA-modifying enzymes identified as candidate positive regulators of exRNAs, we chose to focus on the pseudouridine (Ψ) synthase PUS1^64^, as it was among the top 100 most significant positive regulators (**Fig. 2D, G; Table S2**) and because genetic evidence has implicated Ψ in RNA mobility in *Arabidopsis thaliana* pollen^74^.

To validate the results of the screen, we utilized an orthogonal, pooled CRISPR interference approach (CRISPRi)^75^, whereby we knocked down genes of interest by repressing their transcription with dCas9-KRAB guided by sgRNAs different from those used for the primary screen (**Fig. S3A**). CRISPRi of *Pus1* in CADs suppressed the secretion of sgRNAs significantly (*p =* 0.01) and to a larger extent than the positive control *Sdcbp* (**Fig. S3B**), confirming its involvement in exRNA export.

To determine the contribution of PUS1 to the export of endogenous RNAs, we profiled intra- and extra-cellular RNAs in wild type CADs and two independent *Pus1^-/-^* clones (**Fig. 3A**). Loss of PUS1 resulted in a significantly (adjusted *p* < 0.1) altered profile of 560 exRNAs, skewed more than 2-fold toward reduced export (390 reduced vs. 170 increased exRNAs; **Fig. 3B**). Changes in exRNA abundance were not secondary to altered levels of the corresponding transcripts inside the cells, as shown by a negligible overlap in the affected RNA subsets (**Fig. 3C; S3C–E**). The export of miRNAs, snoRNAs, and both mitochondrial and nuclear tRNAs was significantly impaired in the absence of PUS1 compared to wild type (**Fig. 3D**). Further inspection of tRNA-mapping reads revealed that full-length and 5’ and 3’ tRNA-derived fragments (tDRs; **Fig. 3E**) were exported less efficiently into EVs in absence of PUS1, with the full-length transcripts being most strongly affected (**Fig. 3F**).

**Figure 3.**
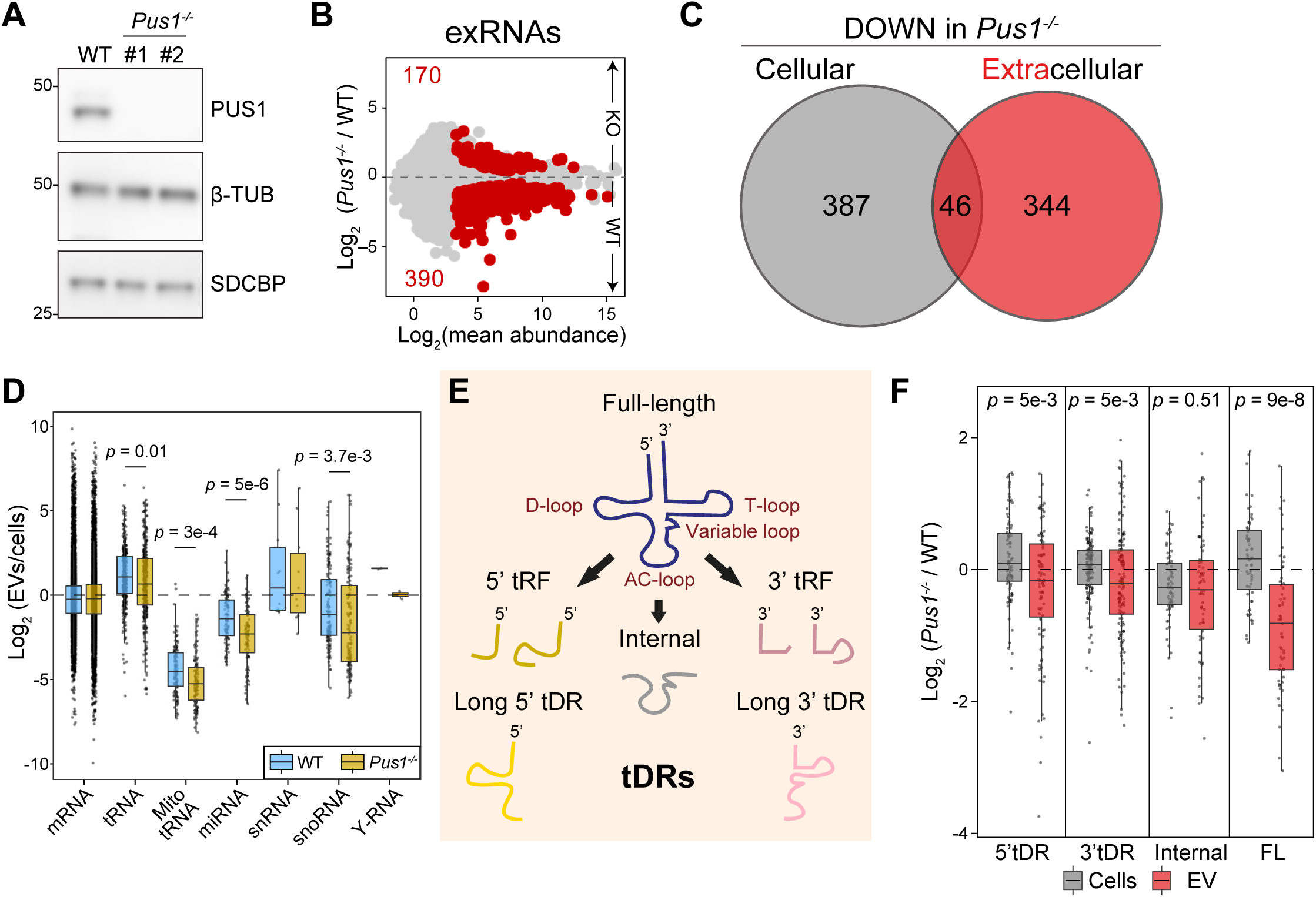
PUS1 facilitates the export of select exRNAs. (A) Western blot probing for PUS1, SDCBP, and β-TUB (control) in wild type (WT) and 2 different *Pus1*^-/-^ CAD clones. (B) Differences in exRNA abundance between wild type and *Pus1*^-/-^, as measured by LIDAR. Red, transcripts showing significant difference (*n* ≥ 3, adjusted *p* < 0.1). (C) Overlap between RNAs less abundant (“DOWN”) in *Pus1*^-/-^ cells compared to WT, either inside the cells (gray, “cellular”) or inside EVs (red, “extracellular”). (D) Ratios of RNA abundance in EVs and inside cells comparing wild type (blue) and *Pus1*^-/-^ (gold) CADs. Average and distributions are shown by the boxes and individual transcripts are shown in points. RNAs were divided in the indicated classes and the *p* values were obtained with Student’s *t* tests. Data from ≥ 3 biological replicates. (E) Scheme showing the different types of known tRNA-derived RNAs (tDRs). (F) Changes in full-length tRNAs or tDRs between wild type and *Pus1*^-/-^ in cells (grey) or EVs (red). *P* values are from a Wilcoxon rank sum test. Data from ≥ 3 biological replicates.

We conclude that PUS1 is necessary for the secretion of certain RNAs, including miRNAs, snoRNAs, and tRNAs, in CADs.

### Pseudouridine stimulates RNA secretion

Based on our observation that PUS1 is required for the release of a subset of exRNAs, we hypothesized that its catalytic product, Ψ, might guide certain RNAs to be incorporated into EVs, facilitating their export into the extracellular space.

To determine if exRNAs contain Ψ, we incorporated the Ψ-bisulfite adduct production step from BID-seq^76^ into our high-sensitivity LIDAR approach^49^. This allowed us to identify Ψ through the appearance of microdeletions in the sequencing reads (**Fig. 4A**), as demonstrated on a synthetic RNA (**Fig. S4A**) and within endogenous 18S rRNA, which contains several known Ψ residues (**Fig. S4B**)^76^. With this approach, we detected 1,111 Ψ sites on 466 distinct exRNAs from CADs (**Fig. S4C–D**). As expected, most Ψ sites were within tRNAs both inside and outside the cells, but RNAs from other classes also contained this modification (**Fig. 4B**), consistent with existing observations^77^. The majority of exRNAs with detectable Ψ also carried this modification inside the cells (**Fig. S4C**), in most cases at the same exact sites (**Fig. S4D**); however, we observed higher Ψ levels on exRNAs compared to cellular RNAs (**Fig. 4C**), suggesting an association of the mark with their extracellular export. We observed a significant (*p* < 10^-15^) enrichment for pseudouridylated RNAs also in EVs isolated from rat primary neuronal cultures via size-exclusion chromatography (**Fig. 4D**), indicating that the role for Ψ in shaping the exRNA population is conserved across neurons from different species and not a peculiarity of transformed CAD cells.

**Figure 4.**
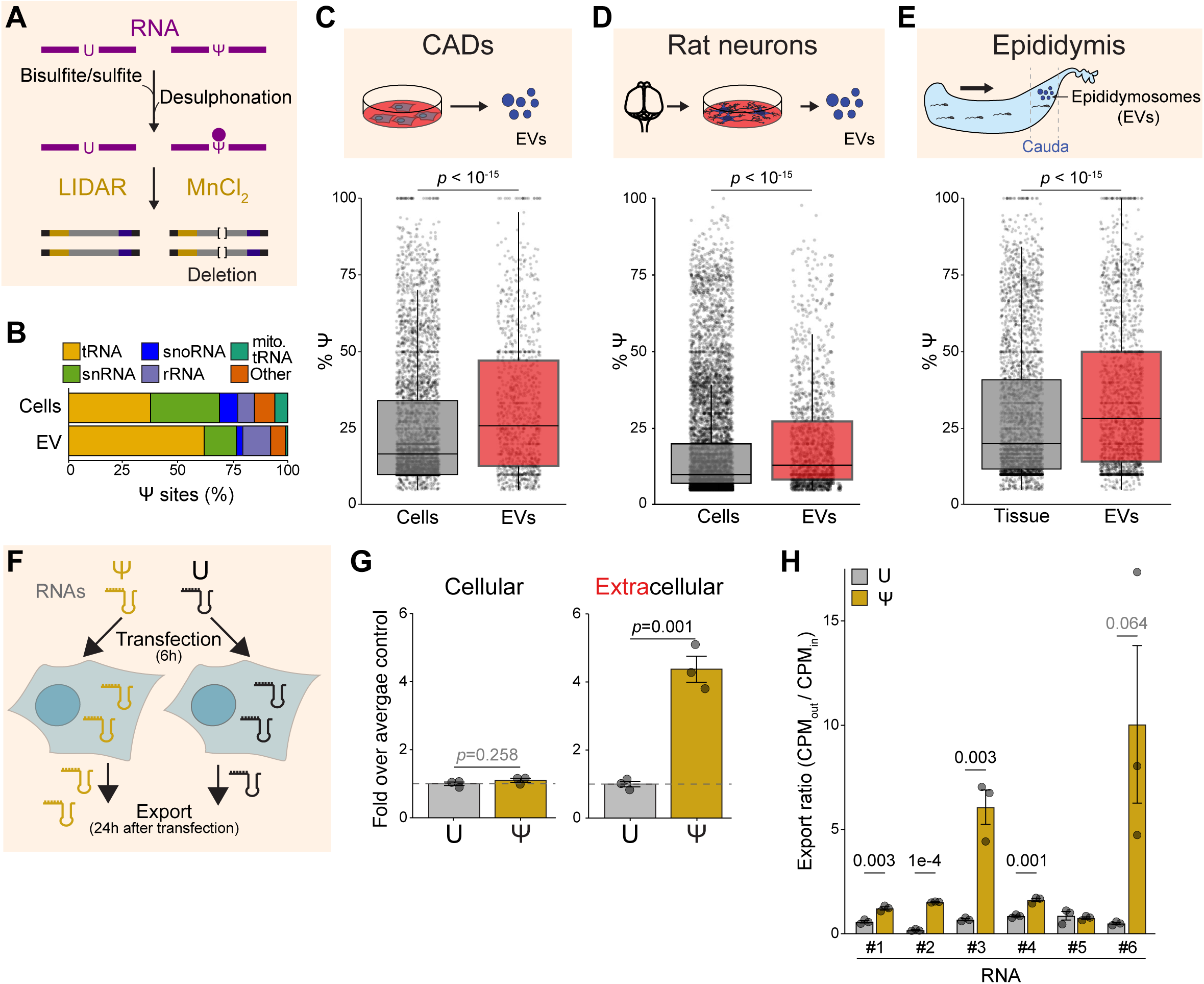
Ψ selects RNAs for secretion. (A) Scheme of the Ψ sequencing approach. Bisulfite-induced deletions were detected with a modified LIDAR protocol which utilizes MnCl_2_ during reverse transcription. (B) Distribution of Ψ-modified RNAs among different classes in CADs and EVs. Data from 2 biological replicates. (C) Ψ levels, inferred by the % of reads containing a deletion at the corresponding sites, for CAD cellular RNAs (gray) and exRNAs (red). *P* value is from a Wilcoxon rank sum test. (D) Same as (C) but for rat primary neurons and corresponding EVs purified by size-exclusion chromatography. (E) Top: Schematic representation of the mouse epididymis, highlighting the region of tissue and EVs used for our analysis (cauda). The direction of sperm movement is indicated by the arrow. Bottom: Same as (C) but for cauda tissue and corresponding EVs. (F) Scheme of the sufficiency experiments shown in (G) and (H). CADs were transfected with RNAs containing only Ψ or U at uridine positions for 6 hours. Their abundance inside and outside cells was measured 24 hours later by RT-qPCR (G) or sequencing (H). (G) Abundance of U- (gray) or Ψ-containing (yellow) RNAs 24 hours after transfection inside and outside CADs, measured by RT-qPCR. *P* values are from Student’s *t* tests. (H) Export ratio (CPM_cells_ / CPM_medium_) for U- (gray) or Ψ-containing (yellow) RNAs, 24 hours after transfection into CADs. Six distinct sequences were tested, indicated by the numbers 1–6. *P* values are from Student’s *t* tests.

Next, we sought to determine whether Ψ was also enriched on exRNAs *in vivo* outside the nervous system. One of the most consequential and intensely studied pathways of extracellular RNA exchange in mammals occurs in the male reproductive tract, where RNAs are transferred to the developing sperm via specialized EVs (epididymosomes) in a process necessary for proper sperm function^78^ and implicated in intergenerational inheritance^12^. We isolated RNAs from EVs purified from the cauda section of the mouse epididymis, (**Fig. 4E; S4E**) and identified 1,976 Ψ sites on mapping to 846 exRNAs. As observed in CADs and primary rat neurons, cauda exRNAs also contained higher levels of Ψ compared to their intracellular counterparts (**Fig. 4E**), demonstrating that Ψ is enriched on exRNAs not only *in vitro* in neurons (see **Fig 4C–D**), but also in other tissues *in vivo*.

Given the enrichment of Ψ on exRNAs, we wondered if Ψ is sufficient to induce RNA secretion. We *in vitro* transcribed 102 bp-long RNAs replacing all U’s with Ψ. We transfected these Ψ-containing RNA into CADs and measured their appearance in the extracellular space by RT-qPCR after 24 hours (**Fig. 4F**). RNA containing Ψ was exported ∼4 times more efficiently compared to the U-containing control, despite their levels being the same inside the cells (**Fig. 4G**). We tested additional RNAs, differing in sequence at their 5’ (**Table S5**) and for most of them (4 out of 6 sequences tested) the presence of Ψ resulted in significantly increased abundance outside the cells, with a fifth one (RNA #6) also showing a clear trend toward increased export (**Fig. 4H**).

Our results strongly suggest that Ψ functions to stimulate export of at least certain subsets of exRNAs in various cell types, both *in vitro* and *in vivo*.

### PUS1 regulates exRNA export via site-specific pseudouridylation

Our genetic evidence linking PUS1 to exRNAs (**Fig. 2–3**) and sequencing and gain-of-function experiments on Ψ (**Fig. 4**) support the hypothesis that conversion of U to Ψ by PUS1 facilitates the secretion of select RNAs. To test this hypothesis, we sought to determine whether the loss of PUS1-dependent Ψ in CADs would cause decreased export of specific exRNAs. Because Ψ can also be deposited by other enzymes, we first identified uridines specifically modified by PUS1, by comparing bisulfite-induced deletions in exRNAs from *Pus1^-/-^* vs. wild type CADs. We found 479 PUS1-sensitive Ψ sites on 270 transcripts, accounting for 43% of the total Ψ sites on exRNAs (**Fig. 5A**). Decreased Ψ levels on these PUS1 targets did not affect their abundance inside the cells compared to RNAs containing Ψ residues unaffected by PUS1 loss (**Fig. 5B**, left). However, the abundance of PUS1 targets in the EVs from *Pus1^-/-^* CADs was significantly decreased (**Fig. 5B**, right) (*p* = 0.008) suggesting that PUS1-mediated deposition of Ψ was required for their efficient export.

**Figure 5.**
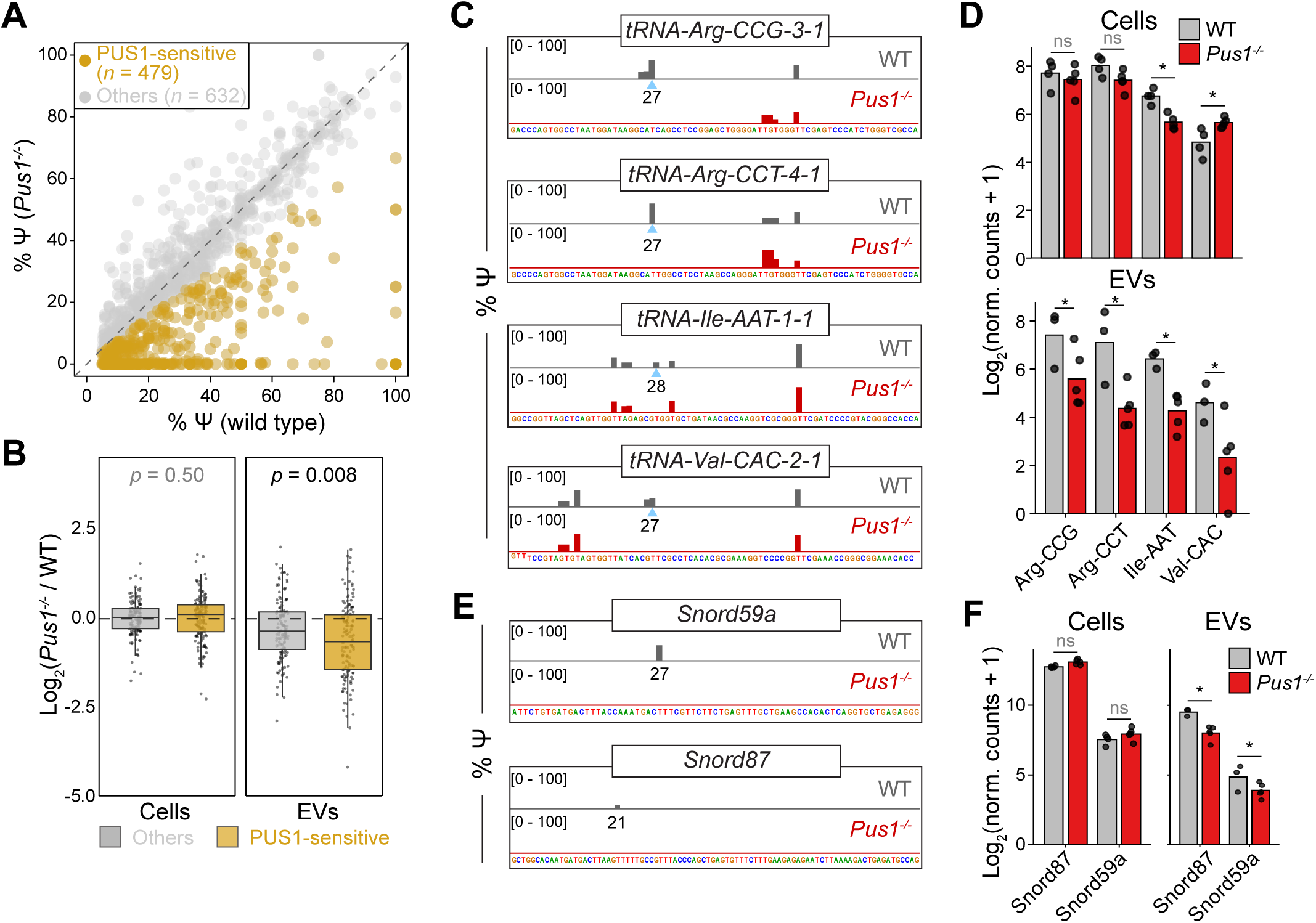
PUS1 marks RNAs for extracellular export. (A) Levels of Ψ modification (% of total reads) at individual exRNA sites in wild type (*x* axis) vs. *Pus1*^-/-^ (*y* axis) CAD EVs. PUS1-sensitive sites, where Ψ was reduced in *Pus1^-/-^* EVs by at least 25%, are in gold. Other Ψ sites are in gray. Data from 2 biological replicates. (B) Ratio of the abundance of Ψ-containing RNAs in *Pus1*^-/-^ CAD vs. WT cells (left) or EVs (right). The RNAs were classified as PUS1-independent (gray, “others”) and PUS1-sensitive (gold). Only RNAs with at least one Ψ site detected in the EV fraction were considered. *P* values are from Wilcoxon rank sum tests. (C) % of molecules containing Ψ at the indicated bases according to bisulfite-LIDAR for four example tRNAs in EVs from wild type (WT) and *Pus1*^-/-^ CAD. (D) Abundance (log_2_-converted, DEseq2 normalized counts) for the full-length tRNAs in (C) in wild type (gray) or *Pus1*^-/-^ (red) EVs. Asterisks indicate *p* < 0.05 (calculated by DEseq2). (E–F) Same as (C) and (D) but for the two indicated snoRNAs.

The majority of exRNAs carrying PUS1-dependent Ψ were full-length tRNAs (**Fig. S5A–B**), which were targeted by PUS1 specifically at position 27 and 28 of the anticodon stem (**Fig. S5C–E**), as previously reported in other cell types^77^. Of the 8 full-length tRNAs that were significantly depleted in EVs from *Pus1^-/-^* CADs, all but one (Fisher’s test *p =* 0.006) contained a PUS1-dependent Ψ at position 27 or 28 (**Fig. S5F**), as exemplified for *tRNA-Arg-CCG-3-1*, *tRNA-Arg-CCT-4-1, tRNA-Ile-AAT-1-1,* and *tRNA-Val-CAC-2-1* (**Fig. 5C–D**)

Not all Ψ sites on these extracellular tRNAs were PUS1 sensitive, as shown by the persistence of Ψ signal at other positions in *Pus1^-/-^*KOs (**Fig. S5D**), most notably base 54, which is modified by PUS10 (**Fig. S5D–E**)^77^. This indicates that the role of Ψ in the export of these exRNAs is context-dependent, and that sequence or structural features of the surrounding RNA (or possibly additional RNA modifications) contribute to its function.

We observed a similar exRNA export defect in *Pus1^-/-^*-derived EVs for snoRNAs (**Fig. S5G**), although in this case only 6 out of 13 snoRNAs containing PUS1-sensitive sites displayed an export defect in the knockout cells, as exemplified by *Snord59a* and *Snord87* (**Fig. 5E–F)**.

Together, these observations support the conclusion that Ψ deposition by PUS1 on specific RNAs is necessary for their efficient extracellular export

### MYL6 mediates the secretion of Ψ-containing RNAs

We hypothesized that one or more RNA-binding protein recognize Ψ-marked RNAs to channel them into secretory pathways. We incubated biotinylated RNA containing Ψ’s or only U’s as control with CAD whole-cell extracts, purified the RNA with streptavidin, and analyzed associated proteins via mass spectrometry. 46 proteins bound specifically to Ψ-containing RNA compared to the U-only control (**Fig. S6A; Table S4**), among which the myosin light chain 6 (MYL6) was the most enriched Ψ-specific binder (**Fig. 6A; Table S4**). MYL6 is an essential light chain component of the non-muscle myosin II complex^79^, and it was previously reported as a putative non-canonical RNA-binding protein^80^.

**Figure 6.**
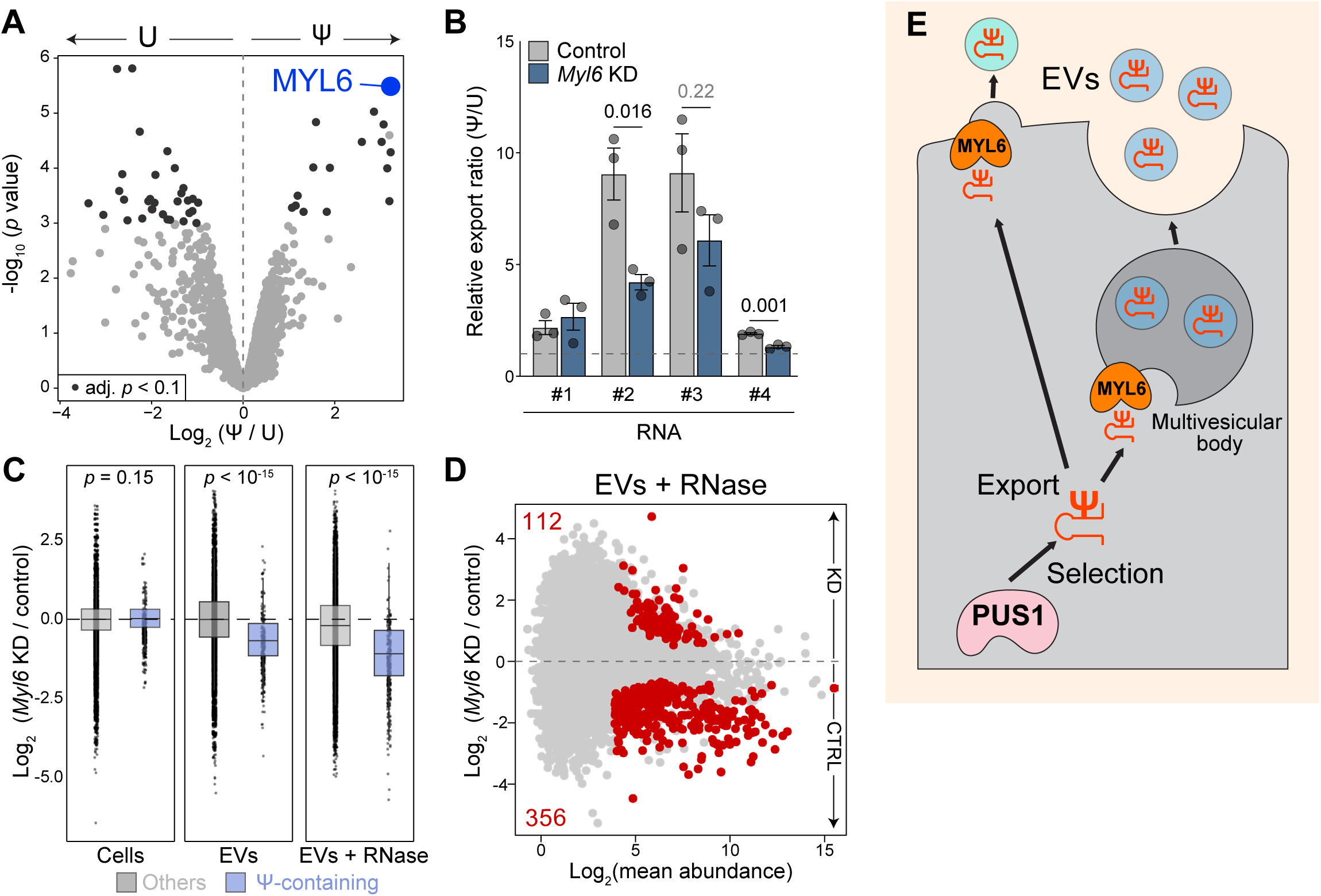
Extracellular export of Ψ-containing RNAs requires MYL6. (A) Comparison of protein abundance in the interactome of U- or Ψ-containing RNA, as measured by mass spectrometry. Dark grey, significantly different proteins (adjusted *p* < 0.1; *n* = 2). (B) Relative export ratio [export ratio(Ψ) / export ratio(U)] between U- and Ψ-containing RNAs in the medium from CADs, treated with siRNAs against *Myl6* and control. The different RNA sequences are identified by the same numbers as in (Fig. 4H). Bars show the mean ± SEM. *P* values are from Student’s *t* tests. (C) Ratio of the abundance of RNAs in *Myl6* knockdown (KD) vs. control in cells (left), EVs (middle) or EVs treated with RNase A/T1 (right). The RNAs were classified as U-containing (gray, “others”) and Ψ-containing (blue). Only RNAs with at least one Ψ site detected in EVs were considered. *P* values are from Wilcoxon rank sum tests. (D) Differences in exRNA abundance (in EVs treated with RNase A/T1) between control and *Myl6* knockdown (KD) as measured by LIDAR. Red, transcripts showing significant difference (adjusted *p* < 0.1; *n* ≥ 2). (E) Proposed model for exRNA secretion. PUS1 marks specific RNAs with Ψ to select them as exRNAs. Once deposited, Ψ provides a binding site for MYL6, which routes RNAs into EVs formed via multivesicular bodies or by direct plasma membrane budding.

Consistent with a role in exRNA trafficking, MYL6 was enriched in the proteome from CAD EVs (**Fig. 1C; Fig. S6B**) and sgRNAs against it were depleted in our screen, although slightly below our significance threshold (*p* = 0.053) (**Fig. S6C**). We knocked down MYL6 in CADs (**Fig. S6D**) and found that Ψ-dependent secretion of 2 out of 4 RNAs was substantially impaired, with a third one also demonstrating a clear trend toward reduction (**Fig. 6B**). To determine whether MYL6 was also involved in the Ψ-dependent export of endogenous exRNAs, we performed LIDAR on intra- and extra-cellular RNAs after MYL6 knockdown. The abundance of intracellular RNAs was largely unaffected (**Fig. S6E**), regardless of their Ψ content (**Fig. 6C**, “cells”); however, the export of Ψ-containing exRNAs in EVs was significantly impaired by MYL6 knockdown (**Fig. 6C–D, Fig. S6F**). This decrease in Ψ-containing exRNAs upon MYL6 knockdown was also observed, and in fact was more pronounced, after RNase treatment of the EV isolate (**Fig. 6C–D**), indicating that the exRNAs affected by MYL6 depletion reside within the EV lumen.

Our results demonstrate that the presence of Ψ—a modification enriched on exRNAs *in vitro* and *in vivo*—is sufficient to stimulate exRNA export, and that, in many cases, Ψ-dependent exRNA sorting in EVs is an active process mediated by the non-canonical RNA-binding protein MYL6.

## DISCUSSION

In this study, we combined biochemical, genetic, and genomic approaches to reveal a role for PUS1 and MYL6 in guiding the extracellular traffic of RNAs via Ψ modification. We propose a model whereby PUS1 converts U to Ψ at defined sites on select RNAs, which are then bound by MYL6 to be packaged into EVs and secreted (**Fig. 6E**). In addition to PUS1 and MYL6, our work points to many new candidate regulators that might play important roles in selecting (RNA-modifying enzymes) and transporting (domesticated retroviral proteins) RNAs from neurons and other cell types into the extracellular space.

### A genome-wide screen for EV and exRNA regulators

We developed a CRISPR/Cas9 pooled genetic screen strategy that linked extracellular phenotypes to cellular genotypes via direct sequencing of secreted sgRNAs (**Fig. 2; S2**). With this design, we could identify not only factors that regulate EV biogenesis in general, but also factors required for the selection, loading, and transport of exRNAs into EVs, and potentially other non-canonical RNA-containing extracellular particles.

Two other screens aimed at separately addressing one or the other process have been published to date. In one case, artificial tethering of sgRNAs though an EV-specific anchor (CD63-dCas9 or CD9-dCas9) was used to create a readout for EV production^56^, which limited the screen to the discovery of factors regulating the biogenesis of these specific EV sub-populations containing CD63 or CD9, and, by design, bypassed the factors required to load RNA into the EVs. In the other, barcoded miRNAs containing the EXO motif^31^ were used as reporters for miRNA secretion, revealing the role of ARHGEF18 in regulating the release of miRNA-containing EVs^81^. In this latter case, only factors involved in movement of this specific subset of exRNAs could be recovered.

While these two studies provided important insights into EV biogenesis and miRNA transport, respectively, our unbiased screen cast a broader net for general regulator of all extracellular particles containing exRNAs, as revealed by the extensive number of “hits” already known to take part in membrane trafficking (**Fig. 2D–E**), and by the large number of candidates involved in RNA biology, such as RNA-modifying enzymes (**Fig. 2G**), that have not been previously linked to RNA secretion.

Importantly, ours was the first genetic screen for an EV phenotype to be conducted on a cell line with neuronal identity, which, in addition to its unique and broader design, likely contributed to our ability to detect several candidates not previously associated with exRNA trafficking and that might be restricted—in expression or function—to the nervous system.

### Routes of exRNA secretion in neurons

While we designed our genetic screen with the goal of uncovering specific regulators of exRNA selection and export, the strategy allowed us to simultaneously identify several factors involved in the biogenesis and trafficking of EVs as “by-products”. The observation that SDCBP was identified as positive regulator (**Fig. S2G–J**) suggests that sgRNAs (and likely other RNAs) taking the extracellular route may be loaded into EVs formed through a variation of the canonical ESCRT pathway, in which SDCBP acts in concert with ALIX to regulate the production of endosomal intraluminal vesicles^59^. This is further supported by the presence of several components of the ESCRTIII complex (CHMP2A, CHMP5, CHMP7) amongst the list of candidate positive regulators (**Fig. 2D**). We also recovered factors potentially involved in alternative modes of exRNA secretion in neuronal cells, such as intraluminal vesicles produced by endosomal membrane curvatures induced by ceramide in lipid rafts^82^, since both SMPD3 (nSmase2, responsible for ceramide synthesis) and FLOT1 (enriched in lipid rafts) appeared amongst our positive regulators of exRNAs (**Table S2**).

In line with the neuronal identity of CADs, but somewhat unexpectedly, we detected synaptic components in our EV preparation (**Fig. 1D; Table S1**) and some of them (e.g.: CXADR, GNAO1, CNR1, ADAM10) were also identified as exRNA regulators in our screen (**Fig. 2E; Table S2**). This suggests that in mammalian neurons, some EVs are produced at the presynaptic membrane, as observed in *Drosophila*^83,84^, indicating the existence of dedicated alternative routes for neuronal exRNAs. Interestingly, fly and mammalian neurons have been shown to package RNAs for secretion in structures resembling viral capsids originating from domesticated retroviral proteins^19,23,85^, whose intercellular exchange appears important for synaptic function, as well as learning and memory^86^.

Among potentially neuron-specific regulators of exRNA trafficking, domesticated retroviral proteins stand out as particularly intriguing, with six recovered in our screen (**Fig. 2F**). One of these, ARC, is a GAG homolog that assembles into a virus-like particle that transports its own mRNA across the plasma membrane in humans and flies^19,20,62^. Several additional genes with homology to the retroviral GAG capsid protein or other retroviral components are under purifying selection in mammalian genomes^60^ and their expression is often high in differentiated neurons^87^ (and also see **Fig. S2K**), hinting at the tantalizing possibilities that brains have hijacked viral machinery for the purpose of RNA-based communication.

### A role for Ψ on exRNAs

Ψ is the most abundant RNA modification found in cells, and it is known for its roles in RNA stability, translation, and splicing^88^. However, the very strong depletion of sgRNAs against PUS1 in our screen, as well as some indirect evidence in other organisms motivated us to explore a potential role in exRNAs. In *Arabidopsis*, Ψ appears to participate to the transfer of small RNAs from pollen to sperm cells^74^. In human colorectal cancer cells, decreased Ψ levels caused by 5-fluorouracil treatment were linked to alteration of extracellular small RNA profiles^89^. In line with these reports, we found Ψ to be: 1) enriched in exRNAs from CAD cells, rat neurons, and mouse epididymis, 2) sufficient for export of most synthetic RNAs tested, and 3) required for the release of endogenous RNA, especially tRNAs and snoRNAs, into the extracellular space.

How does Ψ promote RNA secretion? We propose that the conversion of U to Ψ creates new binding sites for specific proteins that function as Ψ readers. We identified MYL6, part of the non-muscle myosin II complex, as a new Ψ reader and demonstrated its functional involvement in RNA export of synthetic and endogenous exRNAs marked by Ψ **(Fig. 6**). Intriguingly, another cytoskeletal proteins, the actin-binding protein PFN1, has been reported to interact with Ψ in vitro^90^. RNAs containing Ψ might thus “hitchhike” cytoskeletal components to be loaded into vesicles and eventually secreted, a pathway already proposed for other vesicular cargo such as actin-associated proteins^91^. It is worth noting, however, that not all Ψ-containing synthetic and endogenous exRNAs were sensitive to MYL6 depletion. While this could be attributed to experimental limitations (such as incomplete MYL6 knockdown), it is also possible that other Ψ readers are involved, perhaps in an RNA sequence- or structure-dependent fashion.

PUS1 is a conserved eukaryotic standalone pseudouridine synthetase which has been extensively characterized in *S. cerevisiae*^64^, where it plays important roles in tRNA biogenesis^92^. In mammals, PUS1 localizes both to the nuclei and mitochondria^93^. Our work suggests that PUS1-mediated Ψ has a previously unappreciated function: guiding the export of exRNAs. Knockout of PUS1 in neuronal CADs resulted in a significant loss of Ψ from many RNAs and, importantly, in non-overlapping effects on cellular and extracellular RNA pools (**Fig. 3**). PUS1 loss significantly decreased secretion of full-length tRNAs, likely due to the loss of PUS1-dependent Ψ at position 27 and 28 in the anticodon stem. Interestingly, other Ψ sites remain intact, indicating that modification at these other positions may not be sufficient for export (**Fig. 5; S5**) and that the export-related function of Ψ depends on sequence or structural context.

### Chemical modifications for exRNA selection

The most obvious class of candidate positive regulators of exRNAs revealed by our screen were RNA-modifying enzymes (**Fig. 2**). Among these, we chose to focus on PUS1; however, not all pseudouridylated exRNAs depend on PUS1 for their secretion, and many exRNAs do not contain Ψ. We suspect that other RNA modifying enzymes contribute—independently or in concert with PUS1—to shape the exRNA population in neurons and other cell types.

Among these other candidates, we note DTWD1, an enzyme responsible for acp^3^U deposition on tRNAs and possibly other substrates^66^. acp^3^U is an attachment site for N-glycans^94^, and glycosylation of some RNAs seems to be important for their secretion through EVs^95^, suggesting that glycans could serve as alternative exRNA selection marks. Furthermore, we identified ALKBH6 (**Table S2**), a poorly characterized protein predicted to have RNA demethylase activity^96^, as one of the top negative regulators (i.e.: factors that restrict exRNA export). A related RNA demethylates ALKBH5 was recently shown to prevent RNA secretion via m6A removal^97^, and we speculate ALKBH6 may function similarly as a “filter” for exRNA export via modulation of RNA methylation on yet unknown RNA substrates.

Our screen also uncovered hundreds of RNA-binding proteins that may regulate RNA secretion without modifying them enzymatically (**Table S3**). Among these, we identified VAP-A, an important regulator of exRNA export, previously reported to load RNAs into intraluminal vesicles at contact sites between endoplasmic reticulum and multivesicular bodies^98^. We also identified ANXA7, a protein involved in regulating membrane fusion during exocytosis^99^ and recently reported as an RNA binder^100^. Given its enrichment in CAD EVs (**Table S1**) and its reported interaction with ESCRT-III^101^, we postulate it may additional function as an RNA cargo loader during EV formation. On the other hand, we failed to identify some other RNA-binding proteins previously shown to modulate exRNA export, such as ALYREF, FUS, ATG7/12, SYNCRIP, YBX1, and HNRNPA2B1^27,29–33^. This could be due to ineffectiveness of the corresponding sgRNAs, or experimental differences in reporter RNA and cell type used. In fact, it seems possible that different cell types would utilize different sets of RNA-binding proteins and RNA modifications to guide the secretion of different classes of RNAs.

### Conclusions and outlook

In conclusion, our screen in neuronal CADs discovered a new pathway for RNA secretion, whereby PUS1 catalyzes the formation of Ψ on select RNAs, marking them for export via EVs, through recognition by MYL6. The genetic identification and biochemical validation of this and other mechanisms governing exRNA selection and transport will enable deeper mechanistic investigation of their downstream biological functions, and, ultimately, help decode the language of RNA-mediated cell-to-cell communication.

## MATERIALS AND METHODS

### Cell culture and conditioned medium collection

CADs (kind gift from S. Berger)^40^ were maintained in DMEM/F-12 medium (Gibco, Cat. No 11320033) supplemented with 10% ES-grade fetal bovine serum (Gibco, Cat. No 16141079), 1% L-Glutamine (Sigma-Aldrich, Cat. No G7513), and 0.5% penicillin/streptomycin solution (Sigma-Aldrich, Cat. No P0781). EV-depleted fetal bovine serum was generated by centrifugation at 100,000 *g* (28,500 rpm) on a SW-32Ti swinging bucket rotor (Beckman Coulter, Cat. No G369650) for 4 h at 4°C, followed by filtration through a 0.22 μm pore size filter (Sigma-Aldrich, Cat. No S2GPU10RE). For EV collection, cells were seeded at 3–15 million per 15-cm dish and washed twice with 1X PBS. Cells were incubated in serum-free medium (same as maintenance medium but without fetal bovine serum) for 48 h (proteomics) or for 24 h (all other analyses) before collecting EVs. For collection of EVs for pooled CRISPR screen (see below), cells were incubated in EV-depleted medium (same as maintenance medium but with EV-depleted fetal bovine serum) for 24 h. To obtain differentiated CADs, cells were maintained in serum-free medium for 5 days.

### Animals

Male FVB/NJ (8–12 weeks old) were derived from breeders obtained from Jackson Laboratory (Strain #:001800) and maintained in-house. All animal procedures were in strict accordance with the guidelines of the Children’s Hospital of Philadelphia and the University of Pennsylvania Institutional Animal Care and Use Committee regulations (CHOP IACUC Protocol #23-001364, UPenn PSOM IACUC Protocol #806911).

Postnatal day 0-1 Sprague-Dawley rats (Charles River, SAS SD Strain Code 400) of both sexes were used for generating primary neuron cultures. Experiments were performed following protocols approved by the University of Pennsylvania Institutional Animal Care and Use Committee (Protocol #: 807365). Rodents were euthanized following the Humane Euthanasia of Laboratory Animals Standard Operating Procedures of the University of Pennsylvania.

### Dissociated cortical cultures from rats

Bilateral cortices were chopped and digested using 10 mg/mL trypsin and 0.5 mg/mL DNAse for 10 min at 37°C. Tissue was carefully dissociated using a P1000 pipette and cells were plated onto Matrigel (Corning, Cat. No 354234) coated plastic bottom 6-well plates. Neurons were grown following our previously validated astrocyte-free culture approach^50^. Neurons were maintained in Neurobasal (Gibco, Cat. No 21103049) supplemented with 2% B-27 (serum free, Gibco, Cat. No 17504044), 1 mM sodium pyruvate, 2 mM glutamine and 2 μM cytosine arabinoside.

### Extracellular vesicles (EV) isolation and nanoparticle tracking analysis

EV isolation in CADs was performed via differential centrifugation, as previously described^42^ with minor modifications. Briefly, conditioned medium was centrifuged at 300 *g* for 5 min at 4°C, followed by centrifugation at 2,000 *g* for 10 min at 4°C and then 10,000 *g* for 30 min at 4°C. Then, the cleared conditioned medium was passed through a 0.22 μm pore size filter (Millipore-Sigma, Cat. No SCGP00525) and centrifuged at 100,000 *g* (28,500 rpm) on a SW-32Ti swinging bucket rotor for 2 h at 4°C. EV pellet was resuspended in 1X PBS and centrifuged again at 100,000 *g* (28,500 rpm) on a SW-32Ti swinging bucket rotor for 2 h at 4°C. The washed EV pellet was then resuspended in 100 µL of 1 X PBS and either used immediately or stored at -80°C.

Collection of EVs from rat neurons was performed when cortical cultures reached maturity after 14 days *in vitro* (DIV). At DIV 14, 50% of culture media was replaced and 30% of fresh media was added every 24 h. After 60 h, culture supernatant was collected and cleared by serial centrifugations as previously described^50^. The resulting supernatant was used for EVs isolation by size exclusion chromatography using qEV1/35mm columns from IZON (Cat. No IC1-35) and following the procedure suggested by the manufacturer and validated in our previous publication^50^.

For nanoparticle tracking analysis (NTA), EVs were diluted in deionized water and analyzed using the ZetaView PMX220 Twin (Patricle Metrix) with the following parameters: laser wavelength = 488 nm; filter wavelength = scatter; shutter = 100; sensitivity = 80. NTA was performed at the extracellular vesicle core at the University of Pennsylvania.

### Epidydimal tissue and EV collection

Mice were sacrificed by cervical dislocation, and epididymal tissue carefully dissected free of fat and connective tissue. The cauda epididymis was separated and immersed in pre-warmed (37°C) Biggers, Whitten, and Whittingham (BWW) medium^102^. Seminal fluid was gently expressed from the tissue. The tissue was removed from the media and flash frozen in a microcentrifuge tube and kept at -80°C until processing for RNA isolation. The released spermatozoa were incubated in BWW medium at 37 °C for 10 min to allow motile sperm to disperse. The suspension was transferred to a fresh microcentrifuge tube and incubated for an additional 10 min at 37 °C to enrich for motile sperm by swim-up. The upper fraction of the suspension was carefully aspirated, leaving ∼50 µL of residual medium behind. Spermatozoa were pelleted by centrifugation (10,000 *g*, 5 min), and the supernatant was centrifuged twice at 10,000 g for 5 min to remove residual tissue and sperm, each time collecting only the supernatant. A final spin at 10,000 *g* for 30 min at 4 °C was performed to further eliminate debris. The resulting supernatant, containing EVs, was subjected to ultracentrifugation (Optima Max TL, Beckman Coulter) at 120,000 *g* for 2 h at 4 °C. The pellet was washed in 1 mL cold PBS and ultracentrifuged again at 120,000 *g* for 2 h at 4 °C. The final epididymosome pellet was resuspended in 30 µL cold PBS, flash frozen in liquid nitrogen, and stored at -80 °C until RNA isolation.

### Cells and EV proteomics

EVs and cells were resuspended in freshly prepared, ice-cold urea lysis buffer (50 mM Tris pH 8, 8M urea) and incubated for 1 h at 4°C. Lysates were then sonicated using Bioruptor (Diagenode) with 5 cycles of 30s on / 30 s off and then cleared by centrifugation at 18,000 *g* for 10 min at 4°C. Extract was quantified using the Micro BCA Protein Assay Kit (Thermo Scientific, Cat. No 23235) and 50 μg were taken for further processing. Proteins were prepared using Protifi S-trap Micro columns per manufacturer’s instructions. Following S-trap elution, peptides were lyophilized, resuspended in 0.1 % formic acid for further cleanup by desalting. Desalted peptides were lyophilized and resuspended in 4% acetonitrile containing 0.1% formic acid for LC-MS analysis. Mass spectra for peptides were obtained with a Dionex Ultimate3000 liquid chromatograph (Thermo Fisher Scientific) coupled to a Thermo Fusion mass spectrometer (Thermo Fisher Scientific). Peptides were loaded onto an in-house silica capillary column (75 µM inner diameter) packed with C18 resin (ReproSil-Pur C18-AQ 2.4-μm resin; Dr. Maisch GmbH, Ammerbuch, Germany). Peptides were eluted with a gradient of 2%–25% B (86 min) followed by a gradient of 25%–50% (32 min) and a hold at 98% B (15 min) (mobile phase A: 0.1% formic acid, mobile phase B: 80% acetonitrile, 0.1% formic acid). Flow rate was 300 nL/min. Data were collected in data-dependent acquisition mode. Full-scan MS settings were as follows: scan range, 300–1,100 (m/z; mass-to-charge ratio); resolution, 60,000; AGC, on; micro scan count, 1; Maximum IT, 54 ms. MS2 settings were as follows: resolution, 30,000; AGC, predicted; micro scan count, 1; Maximum IT, 100 ms; fragmentation was enforced by higher-energy collisional dissociation with normalized collision energy of 29; isolation width, 1.8 m/z. MS/MS spectra were processed using MetaMorpheus version (https://github.com/smith-chem-wisc/MetaMorpheus) using a UniProt *Mus musculus* reference proteome and common contaminants. The following search settings were used: protease = trypsin; search for truncated proteins and proteolysis products = false; maximum missed cleavages = 2; minimum peptide length = 7; maximum peptide length = 60; initiator methionine behavior = variable; fixed modifications = carbamidomethyl on C, carbamidomethyl on U; variable modifications = oxidation on M; max mods per peptide = 1; max modification isoforms = 1,024; precursor mass tolerance = ±10.0000 PPM; product mass tolerance = ±0.0200 absolute; report PSM ambiguity = true. Results were filtered for score (> 5) and FDR (1 %).

### RNA extraction

Cell pellets or purified EVs were resuspended in 1 mL of TriPure (Roche, Cat. No 11667165001) and RNA was extracted following the recommended protocol. DNA was digested with Turbo DNase-I (Invitrogen, Cat. No AM2238) at 37°C for 30 min, RNA was re-purified using TriPure and resuspended in modBTE (10mM Bis-Tris pH 6.7, 0.1 mM EDTA). For RNA extraction directly from cleared conditioned medium, 0.3 M NaOAc (final) and 1 volume of phenol-chloroform-IAA (Ambion, Cat. No AM9732) were added and RNA was purified using the recommended protocol. DNA was digested and RNA was re-purified using TriPure as described above.

For EVs from cultured primary rat neurons, RNA was extracted from pooled peak fractions using phenol-chloroform-IAA as described for cleared conditioned medium.

### RNase digestion

For RNase protection assays, ∼10^9^-10^10^ EVs were incubated with 2 μg of RNase A + 5U of RNase T1 (Thermo Fisher, Cat. No EN0551) in 100ul of PBS for 30 min at 25°C. Where indicated, Triton X-100 was added to a final concentration of 1% v/v. Reactions were stopped by adding 10 μl of murine RNase inhibitor (New England Biolabs, Cat. No M0314S), followed by incubation on ice for 5 min. Then, 1 mL of TriPure (Roche, Cat. No 11667165001) was added and samples were stored at -80°C. RNA was extracted as described above.

### LIDAR, small RNA-seq, and mRNA-seq libraries

LIDAR libraries from 100–250 ng of cellular RNA or 1–50 ng of EV RNA were prepared as described^49^ with some minor modification. LIDAR_RT_primer and LIDAR_TSO_mix were diluted 1:3 for samples where RNA input was < 10 ng. Reverse transcription conditions were changed to the following: 8°C for 10 min, 16°C for 10 min, 25°C for 10 min, 42°C for 80 min, 10 cycles at 50°C for 2 min and 42°C for 2 min, 85°C for 5 min. Lastly, the amount of indexing primers was increased to 10 pmol. LIDAR libraries were analyzed on a 2% agarose gel and quantified using Qubit dsDNA Quantification Assay kit (Invitrogen, Cat. No Q32854).

Small RNA libraries were prepared using the NEBnext small RNA library kit for Illumina (New England Biolabs, Cat. No E7330S), following the standard protocol with the following parameters: 1) 1:2 dilution of 3’SR Adaptor, SR RT Primer, and 5’SR Adaptor, 2) 15 indexing PCR cycles, 3) cleanup of final libraries with QIAgen MinElute PCR purification kit (QIAgen, Cat. No 28004), followed by a second cleanup with 2X SPRI. Libraries were analyzed on 6% non-denaturing polyacrylamide gels.

Conventional mRNA-seq libraries were prepared from 2 μg of total CAD RNA following the protocol described in^49^.

Libraries were sequenced on Illumina NextSeq1000 or NextSeq2000 instruments.

### Genome-wide pooled CRISPR-Cas9 screen

The mouse Brie sgRNA library^103^ in the lentiCRISPR v2 and lentiGuide-Puro backbones were obtained from Addgene (Cat. No 73632 and 73632). The library were amplified in ElectroMAX Stbl4 competent cells (Invitrogen, Cat. No 11635018). To generate lentiviruses, ∼100 million HEK293FT were transfected with 275 μg of amplified library together with 170 μg of psPAX2 packaging plasmid (Addgene, Cat. No 12260) and 96 µg of pMDG.2 envelope plasmid (Addgene, Cat. No 12259) using Lipofectamine 3000 Transfection Reagent (Invitrogen, Cat. No L3000015). 18 h after transfection, the medium was changed to lentivirus_collection_medium [DMEM high glucose (Gibco, Cat. No 11965126) supplemented with 10% ES-grade fetal bovine serum (Gibco, Cat. No 16141079), 1% GlutaMAX (Gibco, Cat. No 35050061), 1% MEM non-essential amino acid solution (Sigma-Aldrich, Cat. No M7145-100ML), 1% sodium pyruvate (Millipore-Sigma, Cat. No S8636), and 1% bovine serum albumin (Sigma-Aldrich, Cat. No A1933). Conditioned medium was collected 48 h later, filtered through a 0.45 μm filter (Millipore-Sigma, Cat. No S2HVU11RE), and concentrated using Amicon Ultra centrifugal filters with 100 kDa MWCO (Millipore-Sigma, Cat. No UFC910008). Concentrated lentiviruses were aliquoted and stored at -80°C. Multiplicity of infection (MOI) in CADs was estimated as described^104^. Briefly, CADs were infected with increasing amounts of lentivirus using 8 µg/mL polybrene (final concentration). 24 h after infection, cells were divided equally into two new wells and one of them was treated with 1 μg/mL puromycin (Invivogen, Cat. No ant-pr-1). 96 h later, cells were counted, and the MOI was determined as the percentage of surviving cells. To test for sgRNA presence in EV preparations, CADs were infected with the Brie lentiGuide-Puro (sgRNA only) libraries at an MOI of 0.43 and ∼1X coverage, selected with 1 μg/ml puromycin for 7 days, and stably transduced cells were frozen in liquid N_2_. RT-qPCR for sgRNA expression was performed using Power SYBR Green RNA-to-C_T_ (Applied Biosystems, Cat. No 4389986) with the sgRNA_1_qPCR primers (**Table S5**).

The genome-wide screen was performed in 2 biological replicates. 55 million CADs were transduced on day 0 with the lentiviral library at an MOI of 0.21 using 8 μg/mL polybrene to result in ∼100-150X sgRNA coverage. 24 h after infection, cells were washed twice with PBS and EV-depleted medium was added. 24 h later (“day 2” throughout the manuscript), EVs and half the cells were collected as described above and resuspended in TriPure. The other half of the cells was re-plated with maintenance media containing 1 μg/mL puromycin. Cells were selected with puromycin for 7 days. Then, cells were washed twice with PBS and EV-depleted medium was added. 24 h later (“day 10” throughout the manuscript), EVs and cells were collected once again and resuspended in TriPure. RNA was extracted from all samples as described. For cell samples, < 200 nts RNAs were enriched using Zymo RNA Clean & Concentrator – 25 (Zymo Research, Cat. No. R1017) following the recommended protocol.

sgRNA-seq libraries were prepared using a modified LIDAR protocol starting from half of the total amount of RNA obtained from cells and EVs. In the case of cells, multiple parallel libraries were prepared to not exceed 1.5 μg of input RNA / library, which were then pooled before sequencing. Briefly, RNAs was diluted to 2.8 μL and mixed with 0.4 μL of 10 μM LIDAR_sgRNA_RT_primer (see **Table S5** for all oligonucleotide sequences). Samples were heated to 70°C for 2 min, then cooled to 4°C (0.5°C/s). Next, 4.8 μL of RT_mix [25 mM Tris-HCl pH 8.3, 20 mM NaCl, 2.5 mM MgCl_2_, 8 mM DTT, 5% PEG-8000, 0.5 mM dNTPs, 1 mM GTP, 0.32 μL of 50 μM LIDAR_TSO mix^49^ or SS3_TSO, 0.5 U/μL murine RNase inhibitor (New England Biolabs, Cat. No M0314S), and 2U/μL Maxima H-minus RT (Thermo Scientific, Cat. No EP0752); concentrations refer to a final volume of 8 μL] were added. Reverse transcription was performed as follows: 42°C for 90 min, 10 cycles at 50°C for 2 min and 42°C for 2 min, 85°C for 5 min. cDNA was pre-ampified by adding 12 μL of KAPA_mix [1X KAPA HiFi HotStart buffer Ready Mix (Roche, Cat. No KK2601), 1 μM LIDAR_preamp_f, and 0.1 μM LIDAR_preamp_r^49^; concentrations refer to a final volume of 20 μL] and by following these PCR cycling conditions: denaturation (95°C for 3 min), 5 x (98°C for 20 s, 70°C for 30 s, 72°C for 30 s), final extension (72°C for 5 min). Pre-amplified libraries were purified using 2.5X SPRI beads (Beckman Coulter, Cat. No B23319) and eluted in 37 μL of TE buffer (10mM Tris-HCl pH 8, 1 mM EDTA). 0.5 μL of each 10 μM custom Nextera indexing primers was added, followed by 12 μL of Q5_mix [1X Q5 buffer, 0.5 mM dNTPs, 0.02 U/µL Q5 High-Fidelity DNA Polymerase (New England Biolabs, Cat. No M0491); concentrations refer to a final volume of 50 μL]. Samples were incubated with the following PCR cycling conditions: denaturation (98°C for 30 s), 15 x (98°C for 10 s, 65°C for 20 s, 72°C for 20 s). Final libraries were purified using 2X SPRI beads (Beckman Coulter, Cat. No B23319) and eluted in 45 μL of TE buffer (10 mM Tris-HCl pH 8, 1 mM EDTA). Libraries were analyzed on a 2% agarose gel and quantified using Qubit dsDNA Quantification Assay kit (Invitrogen, Cat. No Q32854). Libraries were sequenced on Illumina NextSeq500, NextSeq1000, or NextSeq2000 instruments.

### CRISPRi

CRISPRi sgRNAs targeting the promoter region of *Sdcbp*, *Pus1,* and control (see **Table S5** for all oligonucleotide sequences) were cloned in Lenti-(BB)-EF1a-KRAB-dCas9-P2A-BlastR (Addgene, Cat. No 118154) using Golden Gate. To generate lentiviruses, ∼0.5 million HEK293FT in 6-well plates were transfected with 1.5 μg of CRISPRi plasmid together with 1 μg of psPAX2 packaging plasmid (Addgene, Cat. No 12260) and 0.5 µg of pMDG.2 envelope plasmid (Addgene, Cat. No 12259) using Lipofectamine 3000 Transfection Reagent (Invitrogen, Cat. No L3000015). 24 h after transfection, medium was changed to lentivirus_collection_medium. Conditioned medium was collected 48 h later, and cell and debris were removed by centrifugation at 500 *g* for 5 min, followed by 1,250 *g* for 5 min. Cleared conditioned medium containing lentivirus was concentrated to 50 μL using Amicon Ultra Centrifugal Filter with 100 kDa MWCO (Millipore-Sigma, Cat. No UFC5100). To generate stable CRISPRi cells, 50 thousand CADs were transduced with 50 μL of lentiviruses using 8 µg/mL polybrene (final concentration). 48 h after infection, medium was supplemented with with 20 μg/mL blasticidin (Invivogen, Cat. No ant-bl-1) and cells were selected for 4 days. To generate the pool of different CRISPRi cells, approximately equal number of cells from individual knockdowns were mixed and cells were expanded in the presence of 20 μg/mL blasticidin. Cells were washed twice with PBS and EV-depleted medium was added. 24 h later, conditioned medium was collected and cleared at 300 *g* for 5 min at 4°C, followed by centrifugation at 2,000 *g* for 10 min at 4°C, 10,000 *g* for 30 min at 4°C, and concentration using Amicon Ultra Centrifugal Filter with 100 kDa MWCO (Millipore-Sigma, Cat. No UFC5100). RNA was extracted from cells and concentrated conditioned medium as described above. sgRNA-seq libraries were prepared as described for the genome-wide screen using LIDAR_TSO_mix during the reverse transcription. Libraries were sequenced on Illumina NextSeq1000 or NextSeq2000 instruments.

### Generation of Sdcbp-/- and Pus1-/- cells

Single-guide RNAs targeting *Sdcbp* and *Pus1* (see **Table S5** for all oligonucleotide sequences) were cloned in PX459 (Addgene, Cat. No 62988) using Golden Gate. 0.5 million CADs were transfected with 2.5 μg of the targeting plasmids (PX459 backbone) using Lipofectamine 3000 Transfection Reagent (Invitrogen, Cat. No L3000015). 24 h after transfection, medium was supplemented with 2 μg/mL puromycin, and cells were selected for 48 h. Next, cells were seeded in 15-cm dishes at 500–1,000 cells per dish and incubated for 6 days without puromycin. Individual colonies (corresponding to single clonal populations) were picked and transferred to 96-well plates. Clones were expanded and tested by western blot to assess knock-out efficiency. Whole cell extracts were prepared by lysis of cell pellets in RIPA buffer (10 mM Tris-HCl pH 8, 140 mM NaCl, 1 mM EDTA, 0.1% Na-deoxycholate, 1% Triton-X-100, 0.1% SDS, 1X cOmplete EDTA-free Protease Inhibitor), followed by sonication using Bioruptor (Diagenode) (30 s ON/OFF, 5 cycles) and clarification by centrifugation at 18,000 *g* for 10 min at 4°C.The following antibody dilutions were used for western blots: 1:1000 for αSDCBP (Abcam, Cat. No ab19903), 1:500 for αPUS1 (Abcam, Cat. No ab203010), 1:1000 for αSUZ12 (Cell Signaling Technology, Cat. No 3737S), 1:1000 for αTUBB (Abcam, Cat. No ab6046).

### Export of synthetic sgRNAs

sgRNAs containing uridine or pseudouridine (Ψ) were synthesized by *in vitro* transcription. Briefly, individual sgRNAs were PCR amplified to add a sense T7 promoter sequence (**Table S5**). RNA was transcribed from ∼500 ng of template DNA using HiScribe T7 High Yield RNA Synthesis Kit (New England Biolabs, Cat. No E2040L) following the recommended protocol for short RNAs. UTP was substituted with pseudo-UTP (Jena Bioscience, Cat. No NU-1139L) to generate Ψ-containing sgRNAs. DNA was digested with Turbo DNase-I (Invitrogen, Cat. No AM2238) and RNA was purified using TriPure (Roche, Cat. No 11667165001) as described above.

For assessing export with qPCR, 0.2 million CADs were transfected with 1 μg of UTP- or Ψ-containing EGFP_1_sgRNA using Lipofectamine RNAiMAX Transfection Reagent (Invitrogen, Cat. No 13778075). 6 h after transfection, cells were washed with serum-free medium, and then incubated for additional 20 h in serum-free medium. Conditioned medium was collected and cleared at 300 *g* for 5 min at 4°C, followed by centrifugation at 2,000 *g* for 10 min at 4°C and then 10,000 *g* for 30 min at 4°C. RNA was extracted from cells and cleared conditioned medium as described above. RT-qPCR was performed using Power SYBR Green RNA-to-C_T_ (Applied Biosystems, Cat. No 4389986) with the EGFP_1_qPCR primers (**Table S5**). For assessing export by sequencing, the same procedure was followed with some minor modifications. First, two different sgRNA pools were created mixing 3 UTP-containing and 3 Ψ-containing sgRNAs (**Table S5**). Second, 500 ng of sgRNA pools were used to transfect 0.3 million CADs. RNA was extracted from cells and cleared conditioned medium as described above. sgRNA-seq libraries were prepared using the same strategy as in the genome-wide screen. Libraries were sequenced on Illumina NextSeq1000 or NextSeq2000 instruments.

### Bisulfite-modified LIDAR

Bisulfite conversion and desulphonation of RNA was performed following the BID-seq protocol^76^. Briefly, input RNA (200 ng for cellular RNA and 10 ng for exRNAs) was diluted in 8.5 μL of water, 45 μL of freshly prepared BS reagent (2.4 M Na_2_SO_3_ and 0.36 M NaHSO_3_) were added, and samples were incubated at 70°C for 3 h. Next, 75 μL of water, 270 µL of RNA Binding Buffer (Zymo Research, Cat. No. R1013), and 400 μL of 100% EtOH were added, and samples were transferred on a Zymo column (Zymo Research, Cat. No. R1013). After washing with 200 μL of RNA wash buffer, samples were incubated in-column with 200 μL of RNA desulphonation buffer (Zymo Research, Cat. No. R5001-3-40) for 1 h and 15 min at 25°C. Samples were washed with twice with RNA wash buffer (400 μL and 700 μL), then eluted in 6 μL of RNase-free water. Libraries were prepared starting from half of the eluate using the LIDAR protocol described above with a few modifications: 1) initial RNA denaturation was conducted at 72°C for 3 min; 2) MgCl_2_ was substituted with MnCl_2_ in the RT_mix (keeping the same amounts); 3) reverse transcription conditions were changed to 8°C for 10 min, 16°C for 10 min, 25°C for 10 min, 42°C for 80 min, 10 cycles at 50°C for 2 min and 42°C for 2 min, 85°C for 5 min. Libraries from untreated RNA input samples were prepared in parallel.

### RNA affinity chromatography

Biotinylated sgRNAs containing U or Ψ were generated by *in vitro* transcription. Individual sgRNAs were PCR amplified to add a sense T7 promoter sequence (**Table S5**). RNA was transcribed from ∼600 ng of template DNA (1:1 mix of EGFP_1_sgRNA and SCR_1_sgRNA) using HiScribe T7 High Yield RNA Synthesis Kit (New England Biolabs, Cat. No E2040L) following the recommended protocol for short RNAs. In all reactions, ATP was substitute with a 10:1 mix of ATP and N6-(6-Aminohexyl)-ATP-Biotin (Jena Bioscience, Cat. No NU-805-BIO). UTP was substituted with pseudo-UTP (Jena Bioscience, Cat. No NU-1139L) to generate Ψ-containing sgRNAs. DNA was digested with Turbo DNase-I (Invitrogen, Cat. No AM2238) and RNA was purified using TriPure (Roche, Cat. No 11667165001) as described above.

To prepare protein extracts, approximately 50 million CADs were collected and washed with PBS. Cell pellets were resuspended in 1 mL of ice-cold CHAPS lysis buffer [50 mM Tris-Cl pH = 7.5, 150 mM NaCl, 0.5% CHAPS, 0.5 mM DTT, 1 X cOmplete EDTA-free Protease Inhibitor Cocktail (Roche, Cat. No 11873580001)] and cells were lysed via 2 cycles of freezing in dry ice/EtOH bath followed by rapid thawing at 37°C. Extract was clarified by centrifugation at 18,000 *g* for 10 min at 4°C. Protein amounts were quantified using Bio-Rad Protein Assay Dye Reagent Concentrate (Bio-Rad, Cat. No 5000006).

50 μg of U- or Ψ-containing biotinylated sgRNAs were mixed with 1 mg of CAD protein extract in CHAPS lysis buffer supplemented with 1.5 mM MgCl_2_ and the samples were rotated for 1 h at room temperature. As a control, RNA was omitted from 1 reaction (“beads only”). Next, 50 µL of Dynabeads MyOne Streptavidin C1 beads (Invitrogen, Cat. No 65001), equilibrated in CHAPS lysis buffer supplemented with 1.5 mM MgCl_2_, were added and samples were rotated for 1 h at 4°C. Beads were washed 3 X with medium salt buffer (50 mM Tris-Cl pH = 7.5, 150 mM NaCl, 1.5 mM MgCl_2_, 0.5 mM DTT, 1 X cOmplete EDTA-free Protease Inhibitor Cocktail) and 3 X with high salt buffer (50 mM Tris-Cl pH = 7.5, 500 mM NaCl, 1.5 mM MgCl_2_, 0.5 mM DTT, 1 X cOmplete EDTA-free Protease Inhibitor Cocktail). Proteins were prepared using Protifi S-trap Micro columns per manufacturer’s instructions. Following S-trap elution, peptides were lyophilized, resuspended in 0.1 % formic acid for further cleanup by desalting. Desalted peptides were lyophilized and resuspended in 4% acetonitrile containing 0.1% formic acid for LC-MS analysis. Mass spectra for peptides were obtained with a Dionex Ultimate3000 liquid chromatograph (Thermo Fisher Scientific) coupled to a Q-Exactive HF mass spectrometer (Thermo Fisher Scientific). Peptides were loaded onto an in-house silica capillary column (75 µM inner diameter) packed with C18 resin (ReproSil-Pur C18-AQ 2.4-μm resin; Dr. Maisch GmbH, Ammerbuch, Germany). Peptides were eluted with a gradient of 2%–25% B (85 min) followed by a gradient of 25%–50% (30 min) and a hold at 98% B (15 min) (mobile phase A: 0.1% formic acid, mobile phase B: 80% acetonitrile, 0.1% formic acid). Flow rate was 300 nL/min. Data were collected in data-independent acquisition (DIA) mode had the following settings: Full scan MS – Scan range, 385–1015 m/z; AGC target, 1e6; Ion inject time, 60 ms; Resolution, 60,000; Isolation width, 630. MS/MS scan – Resolution, 30,000; MS2 AGC target, 1e6, Ion inject time, 60 ms; Isolation width, 24; HCD energy, 30. The reference proteomes used for data processing were the Uniprot Mus musculus reference proteome and a Cas9 FASTA (UniprotID: Q99ZW2). DDA spectra were processed with ProteomeDiscoverer (version 2.4.1.15) and Sequest. False discovery rates (FDR) were estimated using a decoy database search with strict and relaxed target FDRs set to 0.01 and 0.05, respectively. DIA spectra were processed with DIA-NN (version 1.9.2). Outputs were filtered at 0.01 FDR, which were estimated based on in silico-generated spectral libraries generated by DIA-NN.

### *Myl6* knock-down

siRNAs against *Myl6* and TYE 563 Control DsiRNA were obtained from Integrated DNA Technologies (Cat. No mm.Ri.Myl6.13.2, mm.Ri.Myl6.13.4, and TriFECTa RNAi Kit). CADs were transfected with siRNAs using Lipofectamine RNAiMAX using 30-50 pmol of siRNA per million cells. Knockdown efficiency was measured 72 h after transfection via RT-qPCR using Power SYBR Green RNA-to-C_T_ (Applied Biosystems, Cat. No 4389986) with the Myl6_qPCR primers (**Table S5**).

### Proteomics data processing and analysis

Protein with measurable intensities only in cell or only in EV samples were excluded from the plot in **Fig. 1C** and denoted as “Cell-exclusive” or “EV-exclusive” in **Table S1**. Raw intensities were normalized using the `median_normalization` function or `proDA` package^105^. Differential abundance analysis was performed in R using the `limma` package as previously described^106,107^. Adjusted *p* values (FDR) were calculated using the `p.adjust` function (method = ‘fdr’). Gene ontology analysis was performed using DAVID (Cellular component 5)^108,109^, including “EV-exclusive” proteins.

### Processing, mapping, and read counting (LIDAR, BID-LIDAR, small RNA-seq, mRNA-seq)

Adapter trimming, UMI extraction, mapping to the *M. musculus* or *R. norvegicus* genome, deduplication and read counting for LIDAR, BID-LIDAR, and small RNA-seq (NEB) was performed exactly as described in^49^ with scripts available on GitHub (https://github.com/bonasio-lab). For mRNA-seq libraries, paired-end reads were mapped directly using STAR^110^ with the following parameters`: --peOverlapNbasesMin 10 --readMapNumber -1 --clip3pNbases 0 --alignIntronMax 100000 --outSAMtype BAM Unsorted`. Counts per gene were calculated in R using summarizeOverlaps function from GenomicAlignments. RNAs with 0 counts in all wild type samples (cellular or exRNAs) were excluded. Raw counts were batch corrected using `Combat_seq`^111^.

### Processing, mapping and, read counting (sgRNA libraries)

Adapter trimming on single-end reads was performed using `Cutadapt` ^112^ with the following parameters: -`a gttttagagcta -m 6`. UMI extraction was performed using the `process_files_single.sh` script available on GitHub (https://github.com/bonasio-lab) with the following parameters: 8bp (libraries constructed using SS3_TSO) or LIDAR (libraries constructed using LIDAR_TSO_mix). Reads were further trimmed with `Cutadapt` with the following parameter: `-u 1`. Reads were mapped to the reference sgRNA Brie library or the sgRNA sequences used in “Export of synthetic sgRNAs” using `bowtie2`^113^ using standard parameters but specifying `--no-rc`. Aligned reads were deduplicated using `umi_tools`^114^ with the following parameter: `--method unique`. Reads shorter than 15 bp were filtered out using `bedtools`^115^. Read counting was performed from bam files using MAGeCK^116^ with the following parameters: `count.` Raw counts were batch corrected using `Combat_seq`^111^.

### CRISPR screen analysis

Batch corrected raw counts were normalized using MAGeCK with the following parameter: `count –norm-method total`. Differential sgRNA abundance between cell and EVs at day 2 was calculated with DESeq2^117^ with the following parameter: `fitType = “local`”.

To identify candidates, first sgRNAs targeting genes not expressed in CADs (RPKM <1) were excluded. Then, to account for individual sgRNA export efficiency as well as changes in sgRNA representation in cells at day 10 due to fitness effects, we calculated a “predicted” sgRNA abundance in day 10 EVs as follows:

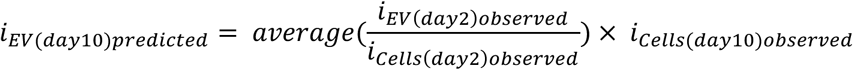

Where *i* is the sgRNA read frequency in the indicated samples. Differences in EV sgRNA predicted and observed abundance at day 10 were calculated using MAGeCK with the following parameter: `test –paired`. GO analysis was performed using DAVID (cellular component, GO level #5)^108,109^.

### Differential RNA abundance analysis (LIDAR)

Differential abundance between wild type cellular and exRNAs, as well as comparisons between wild type and *Pus1^-/-^* mutant CAD, was performed using DESeq2^117^ with the following parameter: `fitType = “local`”. RNAs with an adjusted *p* value < 0.1 were considered differentially enriched/exported.

### Analysis of Ψ with BID-LIDAR

Detection of genomic positions with Ψ was done by identifying positions with deletions induced by BID sequencing compared to a non-bisulfite-induced input. First, reads were mapped using the LIDAR analysis pipeline, as described ab An in-house script using `GenomicRanges`^118^, `BioStrings` (https://bioconductor.org/packages/release/bioc/html/Biostrings.html), and `bedtools`^115^ was used to detect deletions within reads, map putative Ψ sites to the genome reference, and calculate the fraction of reads at each genomic position that contain a deletion. Only positions with at least 5% of reads deleted were considered to harbor a deletion. Positions where deletion occurred 2 X more frequently in BID than input samples were classified as Ψ sites. Position with a BID deletion rate in wild type < 0.75X compared to *Pus1^-/-^* were classified as PUS1-sensitive Ψ sites. These positions were then assigned to the transcripts that contained them to generate a list of modified RNAs.

For tRNA analysis of BID-LIDAR, the position of canonical bases for each tRNA gene were annotated using the strategy described in^49^. These positions were used to identify tRNA containing a U at position 27, which served as candidates for containing a Ψ at this position, and to visualize and quantify reads with deletions at each canonical base.

## DATA AVAILABILITY

Sequencing data is deposited in GEO: GSE320520. Proteomics data is deposited at PRIDE: PXD069544 and PXD069598.

## ACKNOWLEDGMENTS

The authors thank Andy J. Minn for technical advice, and Kenneth Zaret and the Bonasio lab for feedback on the experiments and manuscript. R.B. acknowledges support from the NIH (R35GM153281; DP1NS148061). C.C.C. acknowledges support from the NIH (R35GM151087) This project was supported in part by a Penn Epigenetics Institute Pilot grant to R.B. and A.S.

## AUTHOR CONTRIBUTIONS

Conceptualization: A.S. and R.B.; methodology: A.S., E.J.S., R.B. ; formal analyses: A.S., E.J.S., R.B.; investigation: A.S., T.D.T, E.J.S., L.N.R, J.F.D, J.T., G.E.L., A.A.V., R.L., N.C. ; data curation: A.S., E.J.S., R.B.; writing – original draft: A.S. and R.B.; writing – review & editing: all authors; visualization: A.S., E.J.S., R.B.; supervision: R.B., C.C.C., N.L.C., B.A.G.; project administration: A.S. and R.B.; funding acquisition: R.B.

## SUPPLEMENTARY FIGURE LEGENDS

**Figure S1.**
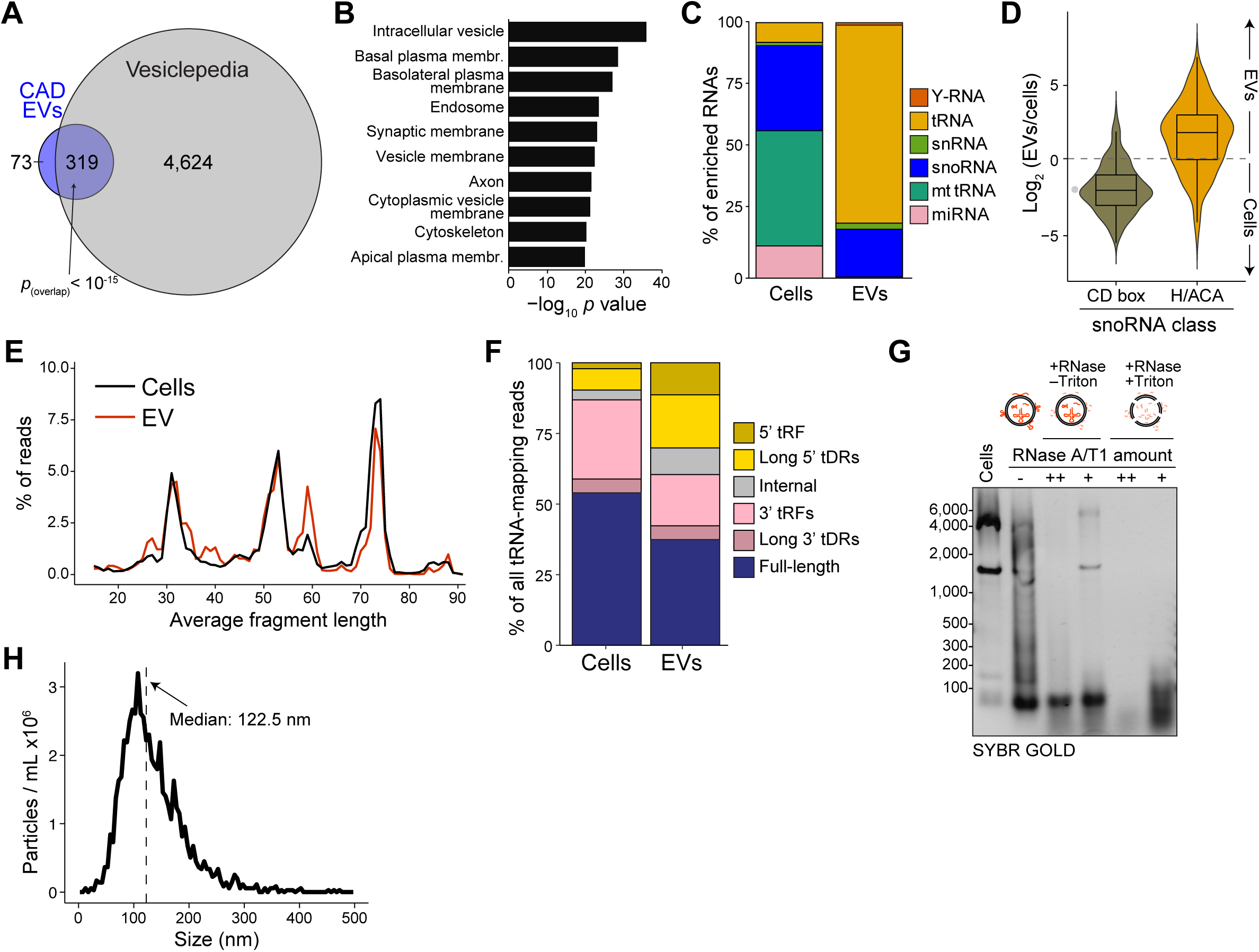
Additional analyses on exRNAs contained in EVs. (A) Overlap between the CAD EV proteome (see Fig. 1C) and proteins in the Vesiclepedia database (Kitti et al., 2024). The *p* value is from a hypergeometric test. (B) Top 9 enriched GO terms (“cellular component” aspect) associated with proteins significantly enriched in EVs from Fig. 1C. (C) RNA class distribution as % of transcripts significantly (adjusted *p* < 0.1) enriched in cells compared to EVs and *vice versa*. (D) Distribution of average (*n* ≥ 3) log-fold-changes between cells and EVs for C/D box and H/ACA box snoRNAs. (E) Average (*n* ≥ 3) insert size distribution of reads mapping to tRNAs in cells and EVs. (F) Average (*n* ≥ 3) tRNA and tDR distribution in cells and EVs. (G) RNAs from EVs subjected to RNase A/T1 digestion in the absence or presence of 1% Triton X-100 were resolved on a 2% agarose-formaldehyde gel. Two amounts of RNaseA/T1 mix were used (++, 2 μg RNase A and 5 U RNase T1 per reaction; +, 0.2 μg RNase A and 0.5 U RNase T1). Cellular RNA (lane 1) and RNA extracted from untreated EVs (lane 2) were loaded as controls. (H) Nanoparticle tracking analysis of EVs purified by size-exclusion chromatography from primary rat neuron cultures. The number of particles per mL is plotted on the *y* axis and their size on the *x* axis. The vertical dashed line indicates the median size.

**Figure S2.**
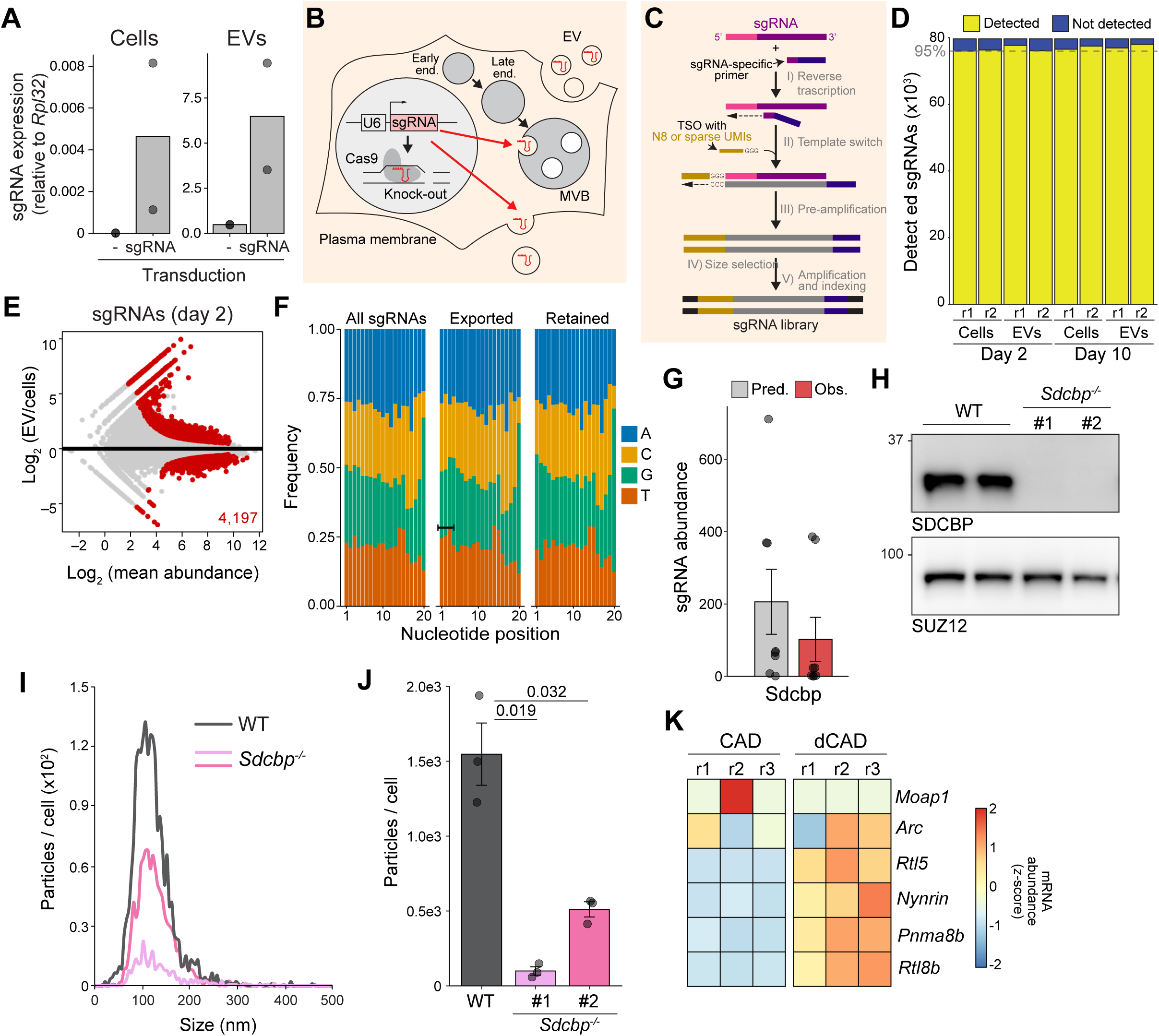
Analyses and validations on the genome-wide screen for exRNAs. (A) RT-qPCR for sgRNA abundance (expressed as ratio to *Rpl32* RNA) in cells and EVs from sgRNA-expressing CAD and controls. (B) Scheme of the logic underpinning the genome-wide screen. A fraction of sgRNAs expressed from a U6 promoter enter endogenous secretion pathways, for example by being loaded into intraluminal vesicles of multivesicular bodies (MVBs, derived from endosomal maturation) or directly by budding from the plasma membrane. This allows for the genotype of the originating cell to be determined via the sgRNA content of the EVs. (C) Scheme of the LIDAR adaptation used to directly sequence the sgRNAs from the EVs. (D) Number of sgRNAs that were detected (yellow) or not (blue) in cells and EV in each screen replicate. Individual samples are shown. Horizontal dashed line indicates the 95% of all sgRNAs included in the library. (E) Comparison of the abundance of all individual sgRNAs inside cells vs. EVs at day 2 after transduction. Red, sgRNAs with significant difference (*n* = 2, adjusted *p* < 0.1). (F) Nucleotide representation at individual CRISPR RNA (crRNA) targeting sequence positions for all, preferentially exported, and preferentially retained sgRNAs. The bracket indicates the first 3 positions, enriched for Us in the exported sgRNAs. (G) Average predicted (gray) and observed (red) values (± SEM) for sgRNAs targeting *Sdcbp.* Circles represent values for the individual sgRNAs. (H) Western blot for SDCBP and SUZ12 (control) in wild type and 2 different *Sdcbp*^-/-^ CAD clones. (I) Nanoparticle tracking analysis of EVs isolated from wild type and *Sdcbp*^-/-^ CADs. The plot shows the number of particles normalized to the number of starting cells on the y axis and their size on the x axis. (J) Average particles/cells (± SEM), secreted by wild type and 2 different *Sdcbp*^-/-^ CAD clones (*n* = 3). Individual replicates are shown. (K) *Z* score-converted mRNA abundance (normalized DEseq2 counts) for the domesticated retroviral genes showed in Fig. 2F, before (“CAD”) and after (“dCAD”) CAD cell differentiation into a more mature neuronal form.

**Figure S3.**
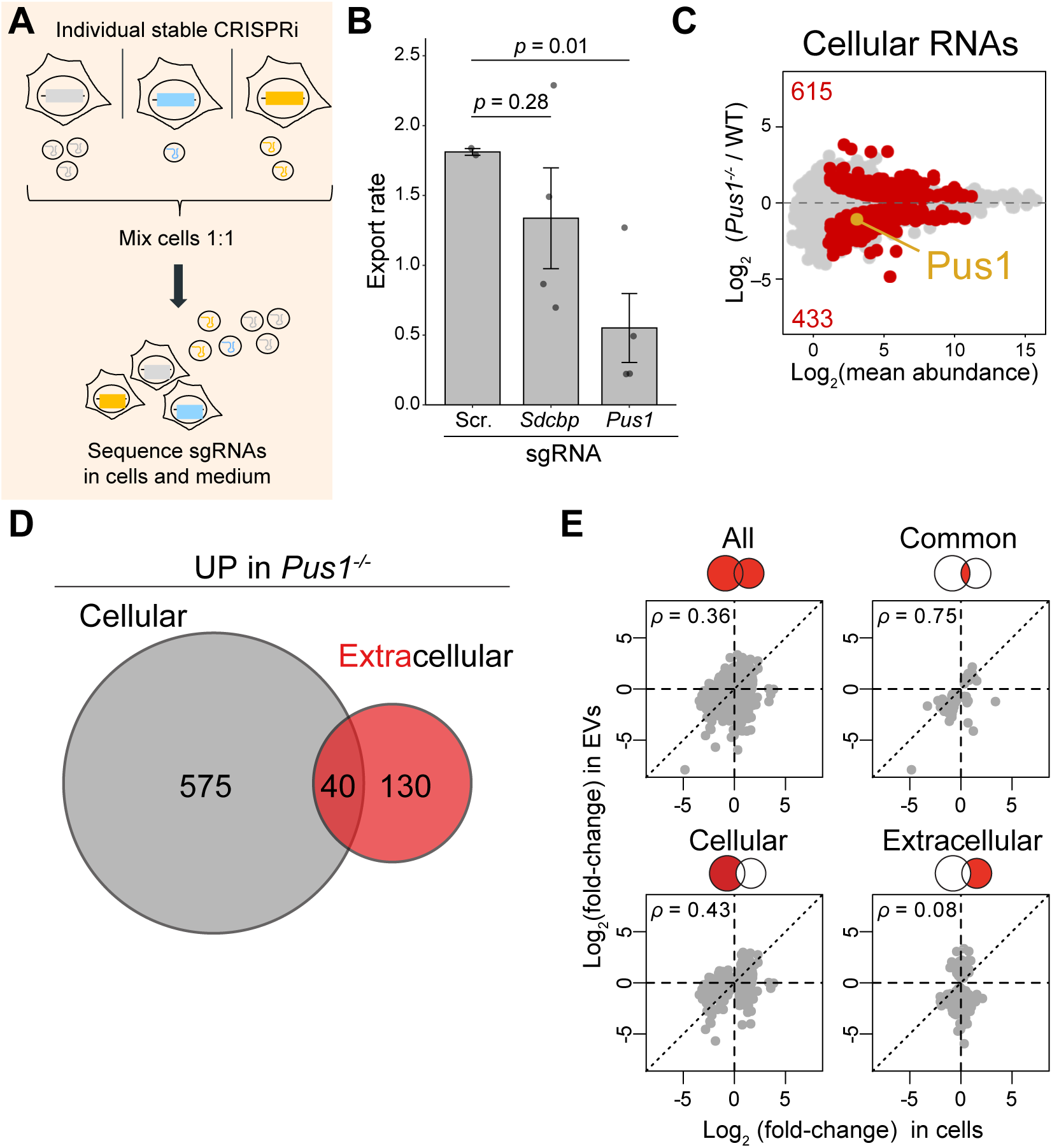
Validation of *Pus1* and analyses on exRNAs from *Pus1^-/-^* cells. (A) Scheme of secondary CRISPRi screen. Stable CADs, transduced with different CRISPRi sgRNAs were mixed in equal numbers. Then, the abundance of sgRNAs in the pooled cells and in the tissue culture medium (extracellular) was measured by LIDAR. (B) Export rate of the indicated CRISPRi sgRNAs, calculated as a ratio of extracellular over intercellular abundance. “Scr.” denotes sgRNA scrambled control. *P* values are from Student’s *t* tests. (C) Differences in cellular RNA abundance between wild type and *Pus1*^-/-^, as measured by LIDAR. Red, transcripts showing significant difference (*n* ≥ 3, adjusted *p* < 0.1). (D) Overlap between RNAs more abundant (“UP”) in *Pus1*^-/-^ cells compared to WT, either inside the cells (gray, “cellular”) or in EVs (red, “extracellular”). (E) Correlations in the change in RNA abundance between wild type and *Pus1*^-/-^ CADs (*x axis*) and EVs (*y* axis) for the RNAs significantly changed (adjusted *p* < 0.1) in either set. Spearman’s correlation values are reported.

**Figure S4.**
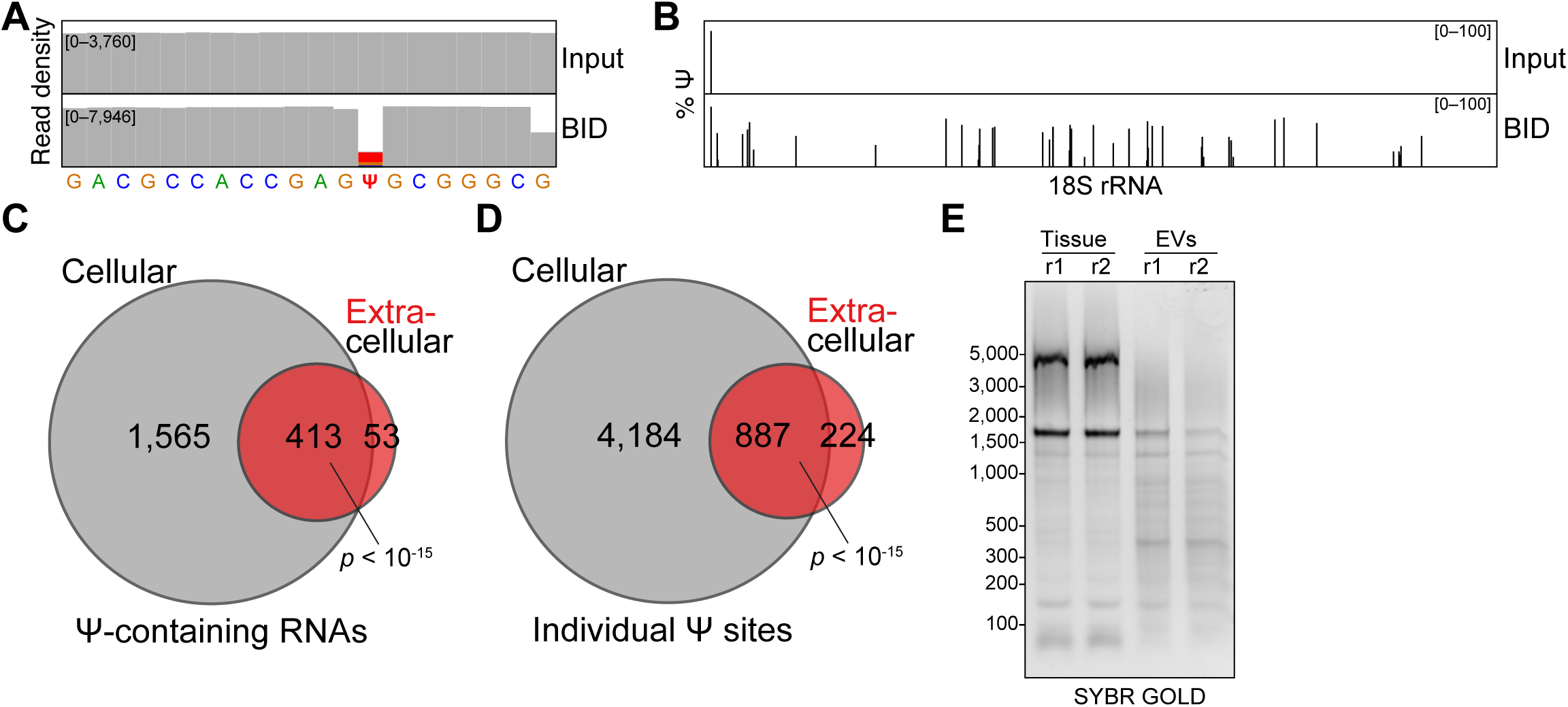
Validation of Ψ sequencing. (A) LIDAR read densities (normalized to counts per millions) for individual nucleotides of an *in vitro*-transcribed Ψ-containing RNA. Top, input; bottom, BID-treated. (B) Inferred % of Ψ based on % of deletion in LIDAR reads mapping to 18s rRNA from CAD RNA. Top, input; bottom, BID-treated. (C–D) Overlap between Ψ-containing RNAs (C) or individual sites (D) identified in CADs and EVs. *P* values are from hypergeometric tests. (E) Denaturing agarose-formaldehyde (2%) electrophoresis of total tissue (cellular) and EV RNA from cauda epididymis (individual replicates are shown).

**Figure S5.**
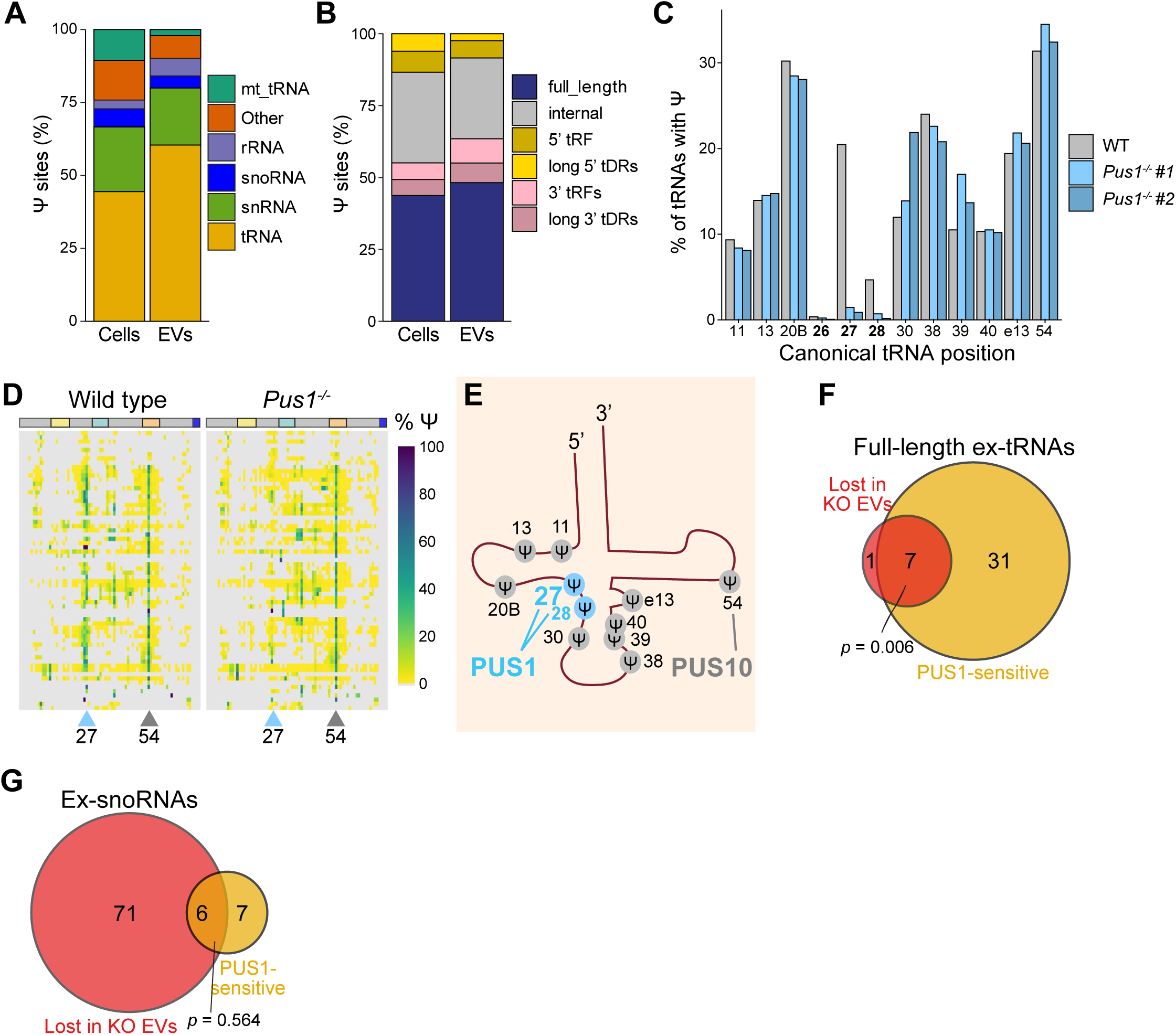
Analyses on exRNAs from *Pus1*^-/-^ CADs. (A) Distribution of transcripts containing PUS1-sensitive Ψ sites in CADs and EVs among the indicated RNA classes. Data from 2 biological replicates. (B) Distribution of reads containing PUS1-sensitive Ψ sites and mapping to tRNAs genes among the indicated types of tRNA-derived species in cells and EVs. (C) % of Ψ modification detected at each canonical tRNA position in wild type or *Pus1^-^*^/-^ CAD EVs. Data from the two different *Pus1^-^*^/-^ clones are shown separately. (D) % of Ψ at every canonical tRNA position (column) for every tRNA iso-acceptor containing a U at position 27 (rows) in EVs from wild type (left) or *Pus1*^-/-^ (right) CADs. (E) Scheme of canonical Ψ sites on tRNAs. In blue, Ψ sites dependent on PUS1. (F) Overlap between significantly reduced full-length tRNAs in *Pus1*^-/-^ (red) and PUS1 full-length tRNA targets (yellow) in EVs. *P* value is from a hypergeometric test. (G) Same as (F) but for snoRNAs. For all panels data were from two biological replicates. In the case of *Pus1^-/-^*the two individual knockout clones were used as combined replicates. For all panels data were from two biological replicates. In the case of *Pus1^-/-^*the two individual knockout clones were used as combined replicates (*n* = 2 per clone).

**Figure S6.**
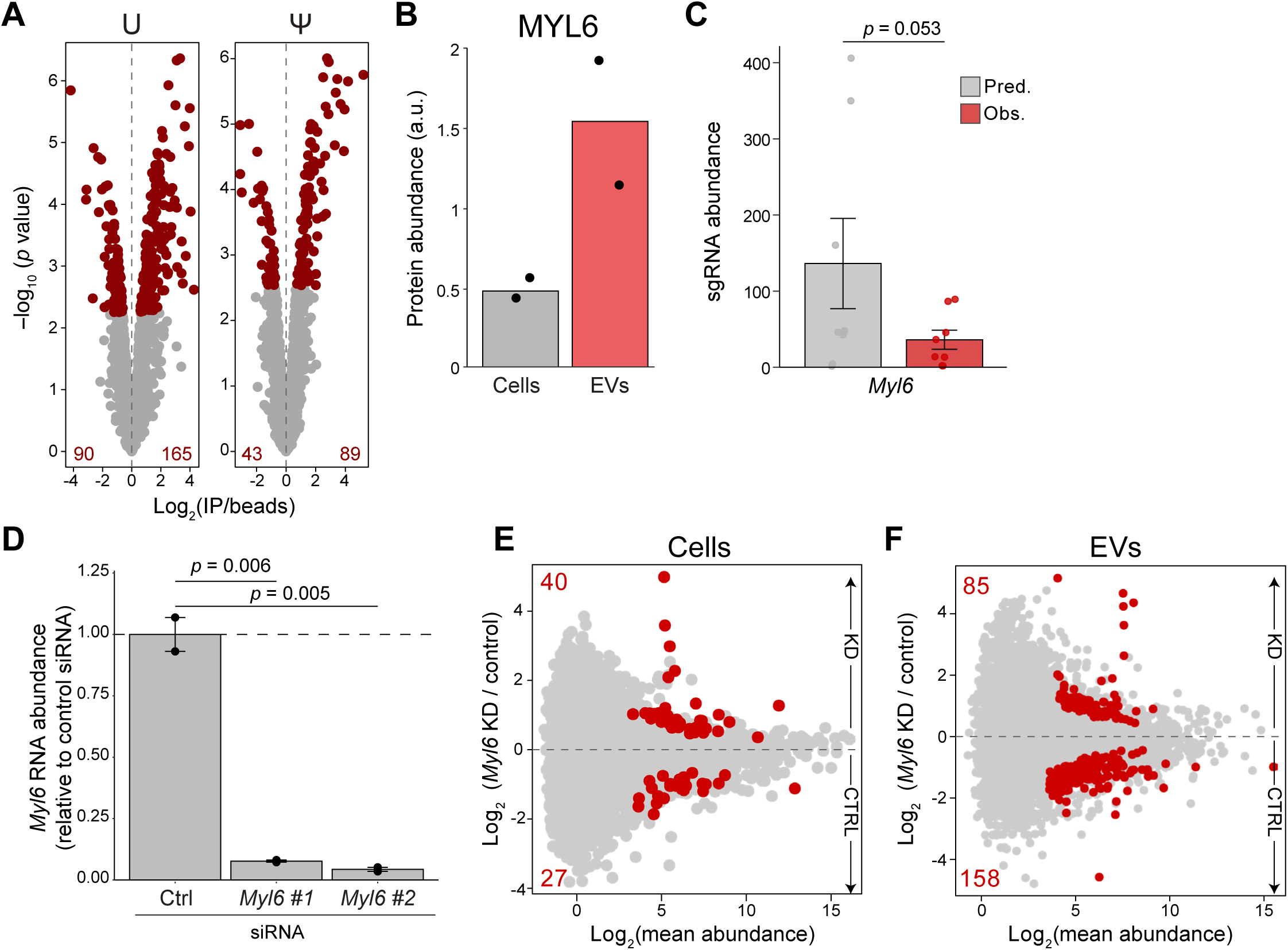
Additional analyses for MYL6. (A) IP/beads values for U- and Ψ-RNAs interactomes measured by mass spectrometry. In red, significantly different proteins (adjusted *p* < 0.1; *n* = 2). (B) MYL6 protein abundance in CADs and EVs. Individual replicates are shown. (C) Average predicted (gray) and observed (red) values (± SEM) for sgRNAs targeting *Myl6.* Circles represent individual sgRNAs. The *p* value is from MaGECK analysis. (D) *Myl6* mRNA abundance relative to control (*Rpl32*). Bars show average ± SEM. Individual replicates are shown. *P* values are from a Student’s t test (E) Differences in intracellular RNA abundance between control and *Myl6* knockdown (KD) as measured by LIDAR. Red, transcripts showing significant difference (adjusted *p* < 0.1; *n* ≥ 2). (F) Same as (E) but for RNAs contained in EVs.

## SUPPLEMENTARY TABLE LEGENDS

**Table S1. Proteomics of CADs and EVs**

**Table S2. Genome-wide screen results**

**Table S3. RNA-binding proteins within positive regulators of exRNA biogenesis**

**Table S4. Proteomics results from U- and Ψ-containing RNA pulldowns**

**Table S5. Oligonucleotides used in this study**

